# Allostery in oligomeric receptor models

**DOI:** 10.1101/330571

**Authors:** Gregory Douglas Conradi Smith

## Abstract

We show how equilibrium binding curves of receptor heterodimers and homodimers can be expressed as rational polynomial functions of the equilibrium binding curves of the constituent monomers, without approximation and without assuming independence of receptor monomers. Using a distinguished spanning tree construction for reduced graph powers, the method properly accounts for thermodynamic constraints and allosteric coupling between receptor monomers.

## Introduction

Guanine nucleotide-binding protein (G protein) coupled receptors (GPCRs) are the largest family of signaling proteins in the mammalian genome and targets for therapeutic drugs (1, 2). When GPCRs are activated by extracellular agonists, they interact with heterotrimeric G proteins to regulate downstream second messenger and protein kinase cascades; notably, cyclic-adenosine monophosphate (cAMP), inositol 1,4,5-triphosphate (IP_3_), and diacylglycerol (DAG).

Equilibrium receptor-occupancy models are used by pharmacologists to quantify changes in ligand affinity and efficacy, and various modes of activation of GPCRs, and to clarify mechanistic hypotheses regarding drug action (3–8). Pharmaceuticals that allosterically modulate GPCRs are of therapeutic interest due to their potential for greater subtype specificity than orthosteric ligands (9, 10). Indeed, allosteric modulators hold promise for treating numerous CNS disorders (11–14).

Evidence for dimerization and oligomerization of GPCRs has been obtained using various experimental methods, including radioligand binding, coimmunoprecipitation, and fluorescence resonance energy transfer microscopy (FRET) (15–17). It is widely believed that dimerization and higher-order complexing (oligomerization) of GPCRs is a common phenomenon that diversifies GPCR signaling and opportunities for pharmacological intervention (18–25).

GPCR dimerization may involve identical receptors (homodimerization), two different subtypes of the same family, or receptors from distantly related families (heterodimerization). Several family C GPCRs exist and function as covalently linked homodimers (e.g., metabotropic glutamate receptors (mGluRs) and calcium-sensing receptors) (26). Some family A GPCRs (e.g., *β*_1_-adenosine and dopamine D_2_ receptors) function as homodimers (27). Some GPCRs are obligate heterodimers (e.g., the GABA_B_ receptor and taste receptors for sweet and umami responses) (28–30). A prototypical GPCR heteromer (composed of receptors from different families) is formed by A_2A_ adenosine receptors and D_2_ dopamine receptors (31–33).

In many of the above examples, physical interactions between monomeric units of GPCR oligomers are known, or suspected to be, important determinants in the mechanism of receptor activation (34–39). Mathematical analysis has provided specific insights into the complexity of allosteric interactions of receptor oligomers (40–45), but a deeper theoretical understanding of oligomeric signaling is needed.

This paper introduces a novel theoretical framework for understanding allostery and thermodynamic constraints in oligomeric receptor models that are composed of any number of identical monomers. The framework allows equilibrium occupancy measures (i.e., binding curves) of receptor homodimers to be expressed in terms of the properties of constituent monomers, without approximation and without assuming independence of receptor monomers. This is achieved by constructing the state-transition diagram of the receptor oligomer, identifying thermodynamic constraints, and constructing a distinguished spanning tree of this graph, as explained below. The mathematics in the main text will be familiar to quantitative pharmacologists. The Supplementary Notes presents results with mathematical rigor in the language of algebraic graph theory (product graphs, cycle bases, and so on).

## Methods

This paper presumes basic understanding of equilibrium receptor-occupancy models as used by the mainstream pharmacological community (see (46) for an overview). In this section we review this methodology and, *en passant*, distinguish two ways that thermodynamic constraints and allosteric parameters arise in receptor models: (1) when the state-transition graph of a receptor includes cycles (as in Fig. 1), and (2) as a property of receptor oligomers that emerges via conformational coupling of constituent monomers.

**Fig. 1.**
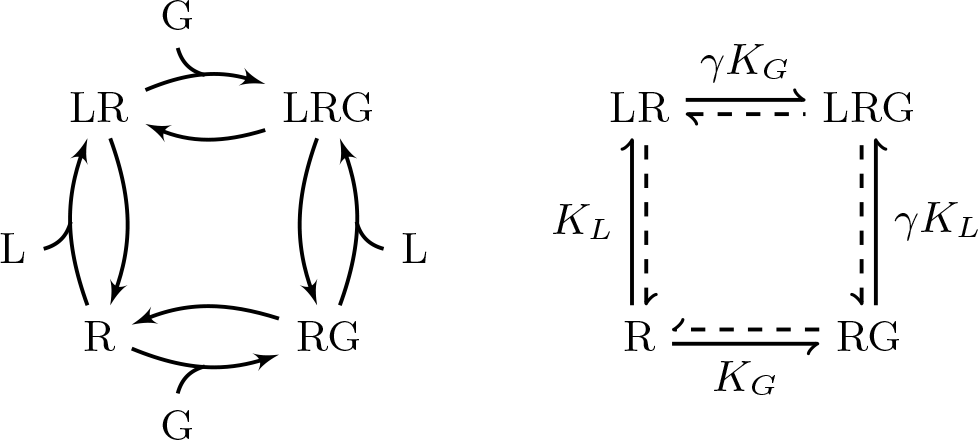
Ternary complex model of a G protein coupled receptor (47–51). For given ligand and G protein concentration ([L] and [G]), there are three free parameters: *K*_*L*_, *K_G_* and *γ*.

### Thermodynamic constraints and allostery

It is well-known that G proteins may modulate ligand affinity at GPCRs (47–51). This phenomenon illustrates important relationships between cycles in the graph representing receptor model topology, thermodynamic constraints on equilibrium model parameters, and allosteric coupling (46, 52).

Consider the ternary complex model of ligand (L), 7-transmembrane receptor (R), and G protein (G) interactions (Fig. 1). As will be familiar to many readers, the ternary complex model hypothesizes distinct binding sites for ligand (orthosteric) and G protein (allosteric), 4 receptor conformations (states), and 4 reversible reactions. Microscopic reversibility requires that the product of the transition rates around the four states of the ternary complex model is the same clockwise as counter-clockwise when ligand and G protein concentrations are independent of receptor state (53). If we consider bimolecular association as the forward reaction, the chemical equilibrium constants are *K*_*L*_ = [LR]/([L][R]), *K*_*G*_ = [RG]/([G][R]), 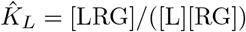, and 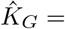 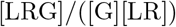, where [L], [R], etc., represent equilibrium concentrations. The cycle in the ternary complex model leads to the thermodynamic constraint 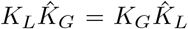 and, consequently, the TCM has 3 (not 4) free equilibrium parameters. To emphasize the *cooperativity* of the two binding processes, one may define an allosteric parameter 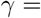 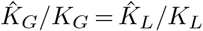. In that case, the receptor model is specified by two equilibrium association constants (*K*_*L*_, *K*_*G*_) and *γ*, the strength of allosteric coupling. The ligand affinity is *K*_*L*_ when G protein is unbound, and *γK*_*L*_ when G protein is bound. Thus, *γ* > 1 specifies a TCM in which G protein binding increases agonist affinity, as observed for *β*_2_-ARs (47–49). Fig. S1 in the Supplementary Notes shows binding curves for the TCM model given by [R]/[R]_*T*_, [LR]/[R]_*T*_, etc., as a function of [L] and [G].

### Cooperativity in receptor dimers

Thermodynamic constraints and allosteric parameters also arise when modeling receptor dimers and the interactions between constituent monomers. To illustrate, consider a monomer with sequential binding reactions,

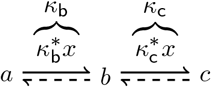

where *κ*_b_ and *κ*_c_ are dimensionless equilibrium constants and *x* is ligand concentration. In this diagram, solid harpoons indicate the forward reaction direction. For example, the reaction labelled *κ*_b_ has *a* as reactant and *b* as product; consequently, increasing *κ*_b_ decreases the occupancy of state *a* and increases the occupancy of state *b*. The diagram introduces helpful notation. The states are labelled in such a way that the reactant comes before the product in dictionary order (*a* to *b* to *c*). The subscript of the equilibrium constants *κ*_b_ and *κ*_c_ are chosen to match the label of the reaction products.

For an isolated monomer, the occupancy measures are given by *π_i_* = *z_i_/z_T_* where *z_T_* = ∑_*i*_ *z_i_*. *z_a_* = 1, *z_b_* = *κ*_b_, and *z_c_* = *κ*_c_*κ*_b_, that is,

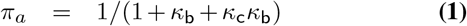

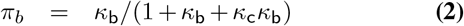

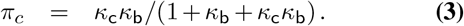

It is convenient to denote this set of rational functions as

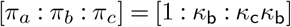

where it is understood that [*x*_1_: *x*_2_: …: *x_n_*] = [*λx*_1_: *λx*_2_: …: *λx_n_*] for any *λ* ≠ 0, and *λ* ≠ 1/ ∑_*i*_*x_n_* gives the normalized probability distribution *π* = (*π*_1_, *π*_2_,…, *π_n_*) where 1 = ∑_*i*_ *π_i_*. Fig. 2 shows representative binding curves given by Eqs. 1–3 as a function of the ligand concentration *x*.

### Heterodimer

A receptor heterodimer model composed of two *distinguishable* monomers with this 3-state topology has 9 states, 12 reversible reactions, 4 thermodynamic constraints, and 12 − 4 = 8 free equilibrium parameters (Fig. 3A). Each monomer contributes 2 parameters, for a total of 4 (*κ*_b_, *κ*_c_ and 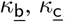). The remaining parameters (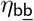, 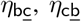 and 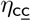) encode the strength of 4 2-way allosteric interactions among the monomers (one for each 4-cycle).

**Fig. 2.**
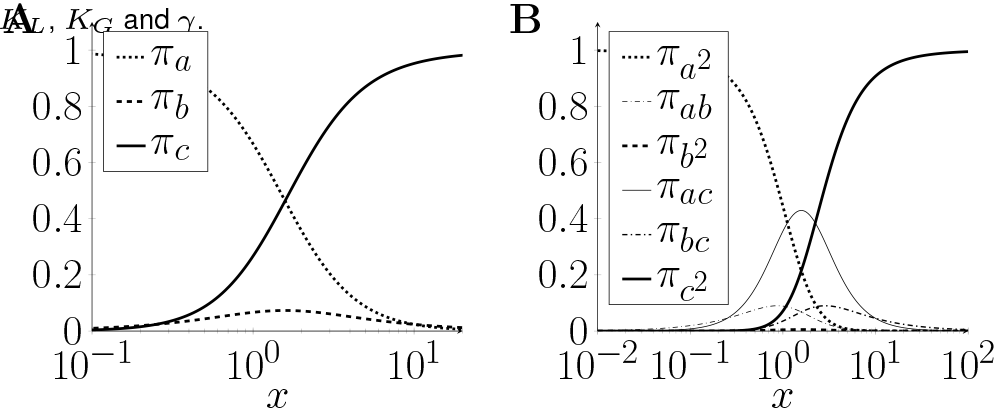
(A) Binding curves for monomer with sequential binding reactions (Eqs. 1–3) with 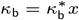 and 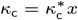, and association constants 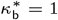 and 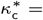 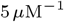. (B) Binding curves for dimer given by Eq. 5 with *η_i_* = 1.

**Fig. 3.**
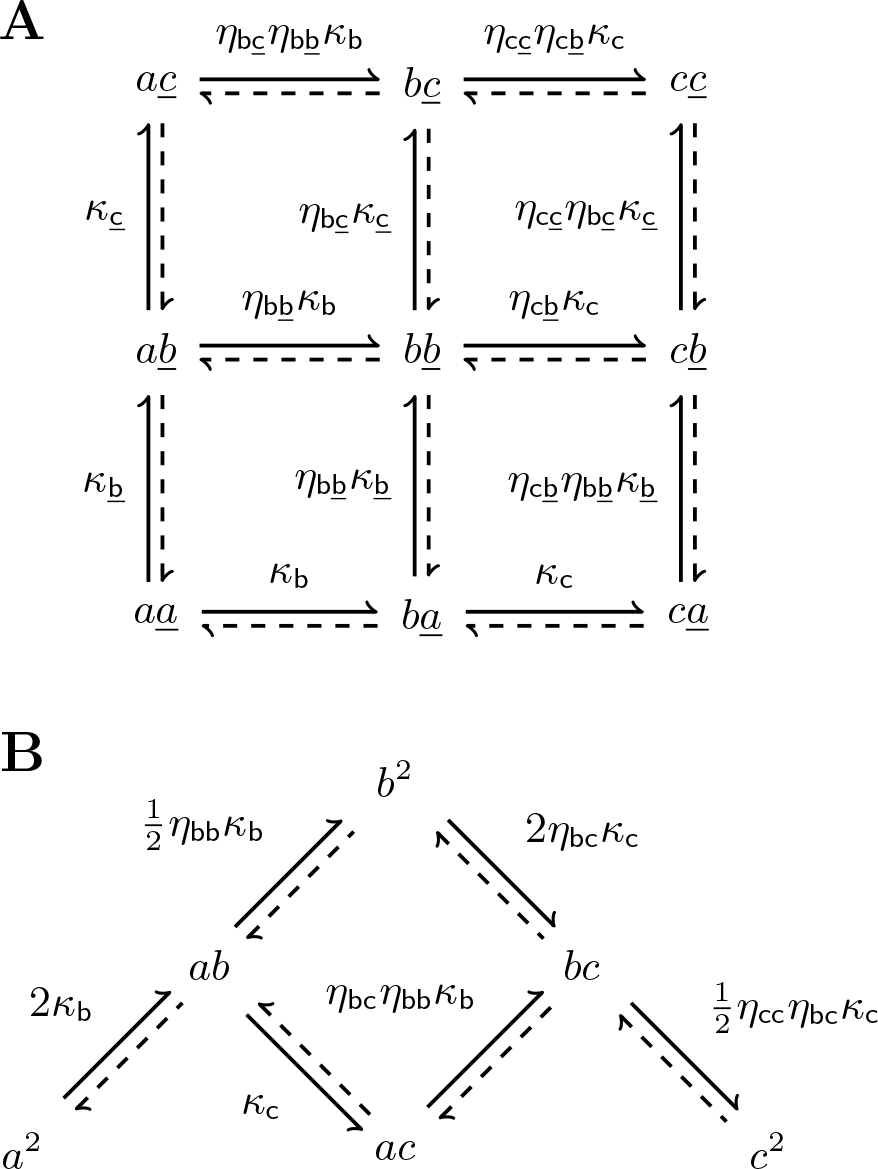
(A) Equilibrium parameters for a heterodimer composed of two 3-state monomers (same topology, different parameters). There are four 4-cycles and four allosteric parameters: 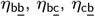 and 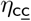. Right: State-transition diagrams for dimer composed of identical and indistinguishable subunits (not necessarily independent) and three allosteric parameters: *η*_bb_, *η*_bc_, and *η*_cc_.

To clarify the meaning of allosteric parameters in Fig. 3, write 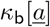 for the equilibrium constant of the *a* ⇌ *b* reaction of first monomer *occurring in the context of the second monomer being in state* 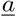, and similarly for 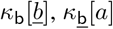, 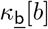.

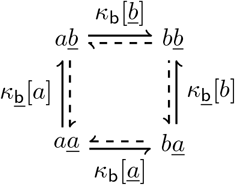

In that case, the allosteric parameter 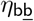 is, by definition,

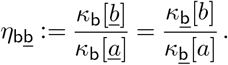

Taking states *a* and 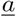 as reference states, we write 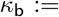 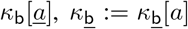. Consequently, 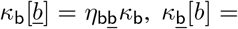, 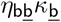, and the equilibrium parameters are

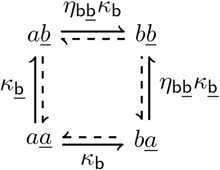

Similar definitions for 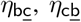 and 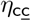 lead to state-transition diagram of Fig. 3. The proportion of dimers in each state,

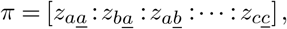

can be ‘read off’ the state-transition diagram, remembering that the equilibrium constants are defined so that *a → b → c* and 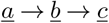 are forward reactions (52). Because 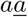 is not a product of a forward reaction, we assign 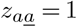. The other *z_i_* are given by the product of equilibrium parameters labeling forward reactions on a path from 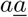 to *i*. For example, to calculate 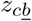, we observe the path 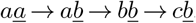, passes in the forward direction through three reactions with equilibrium constants 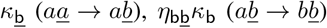, and 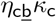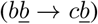; the product gives 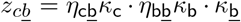. In a similar manner we obtain 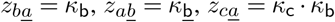, 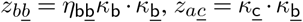,

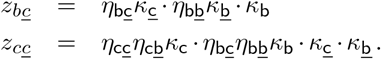

### Homodimer

A receptor homodimer composed of two *indistinguishable* monomers with the same 3-state topology has 6 states, 6 reversible reactions, 1 thermodynamic constraint, and 6−1 = 5 free equilibrium parameters (Fig. 3B). The homodimer state-transition diagram (B) is a contraction of the heterodimer diagram (A) obtained by lumping and renaming states 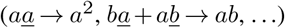 and parameters (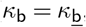, 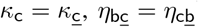). The monomers, being identical, contribute only 2 parameters (*κ*_b_, *κ*_c_). There are only three distinct allosteric parameters for the stength of 2-way interactions, written as *η*_bb_, *η*_bc_ and *η*_cc_. The fraction of dimers in each state,

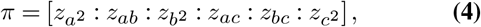

is given by 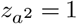, *z_ab_* = 2*κ*_b_, 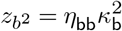, *z_ac_* = 2*κ*_b_*κ*c,

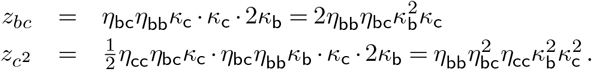

Where the combinatorial coefficient 2 (resp. 1/2) appears as a factor on the transitions out of (resp. into) states *a*^2^, *b*^2^ and *c*^2^.

### Homodimer allostery

Importantly, the above calculation did *not* assume independent monomers. Rather, the dependence of the monomers in the homodimer has been parameterized by the three allosteric parameters *η*_bb_, *η*_bc_ and *η*_cc_. To see this, transform Eq. 4 to an equivalent expression by dividing each term by (1 + *κ*_b_ + *κ*_c_*κ*_b_)^2^ to obtain

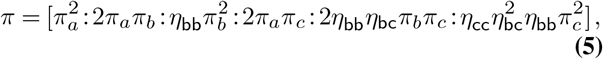

where *π_a_*, *π_b_* and *π_c_* are given by Eqs. 1–3.

Eq. 5 is significant. *Without assuming independence*, we have expressed the occupancy measure for a receptor homodimer in terms of…

- the occupancy measures of a isolated monomer (*π_a_*, *π*_*b*_ and *π_c_* determined by *κ_b_*, and *κ_c_*) and
- the allosteric parameters *η*_bb_, *η*_bc_ and *η*_cc_.

In the absence of allosteric interactions, *η*_*i*_ = 1 and Eq. 5 simplifies as expected: 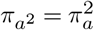, *π_ab_*= 2*π_a_π_b_*, etc.

To illustrate the effect of conformational coupling between monomers of the 6-state dimer (Fig. 3), Fig. 4A plots the fraction of occupied ligand binding sites (4 in the dimer, 2 for each monomer),

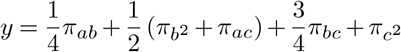

as a function of ligand concentration. In terms of the monomer occupation measures and allosteric parameters, we find

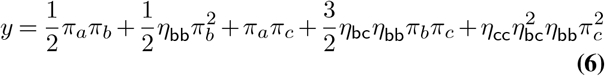

where we have used 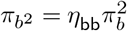, *π_bc_* = 2*η*_bc_*η*_bb_*π_b_π_c_*, etc., obtained by identifying Eqs. 4 and 5. The Hill plots in Fig. 4, B and C, show how the allosteric parameters (*η*_bb_, *η*_bc_ and *η*_cc_) that characterize the interactions between the monomers may lead to cooperativity in the fraction of occupied binding sites. This example is reminiscent of a sequential (as opposed to concerted) model of cooperative oxygen binding in hemoglobin that accounts for the inequivalence of *α* and *β* subunits (54, 55). In this interpretation, the original three-state model is analogous to a *αβ* hemoglobin dimer, and the allosteric parameter *η*_bb_ is the increase in affinity for the second binding event. The 6-state model represents a hemoglobin tetramer, in which *η*_bc_ and *η*_cc_ represent affinity changes resulting from interactions between *αβ* dimers.

**Fig. 4.**
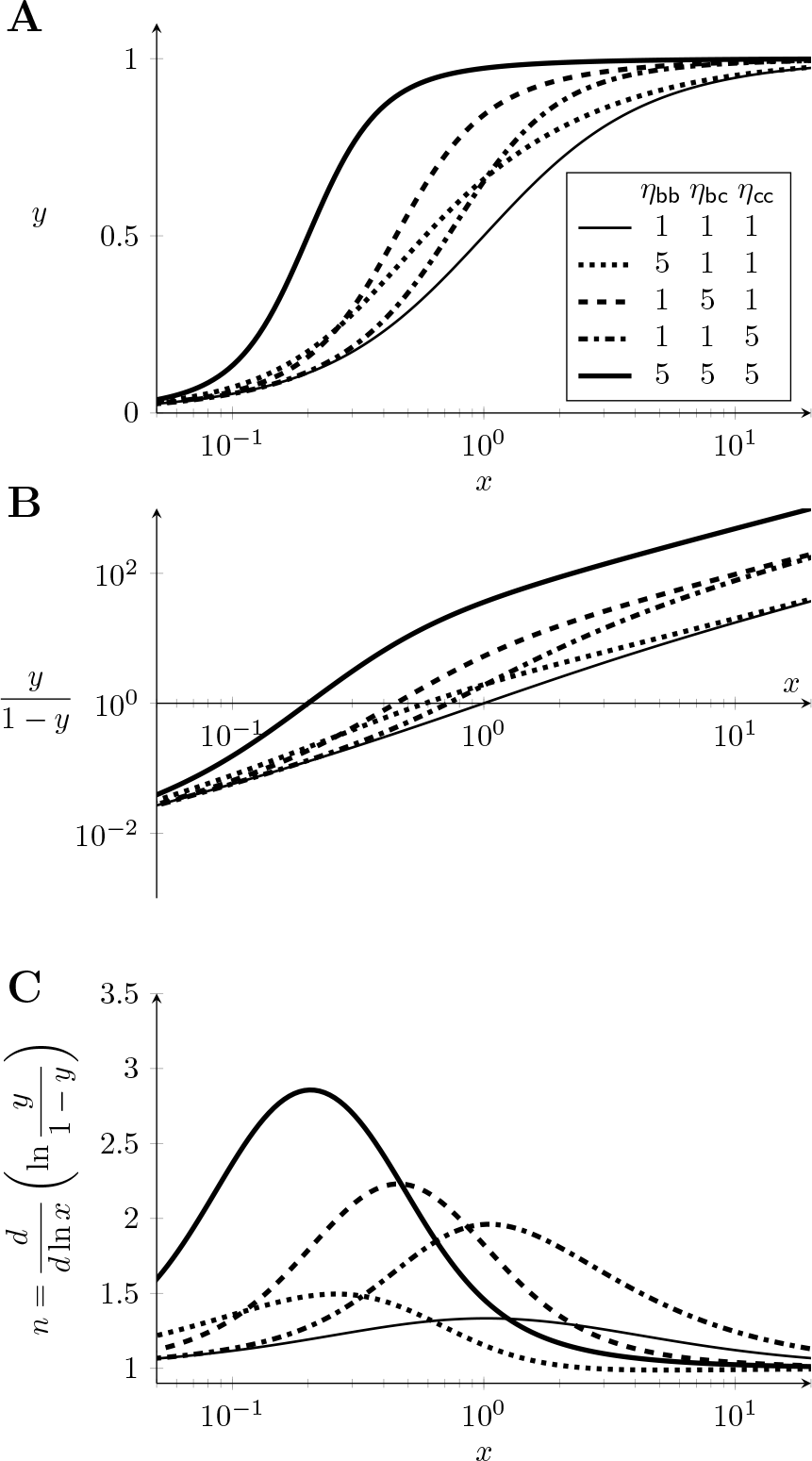
(A) The fraction of occupied ligand binding sites (Eqs. and 6) in the homodimer model (Fig. 3B) for different values of the allosteric parameters *η*_bb_, *η*_bc_ and *η*_cc_. (B,C) Hill plots show that interactions between the monomers may lead to cooperativity.

The remainder of the paper presents a *general theory of allostery in oligomeric receptors composed of any number of identical monomers*. First, we provide a construction of the state-transition diagram of receptor homodimers (and oligomers), for any given monomer topology. Next, we characterize cycles and thermodynamic constraints in receptor oligomers *in terms of the monomer topology* (i.e., without having to construct the state-transition graph of the oligomer). Third, we show how allosteric interactions in receptor oligomers may be systematically enumerated. Fourth, we show how the occupancy measure of a receptor oligomer may always be expressed in terms of the occupancy measures of an isolated monomer and identified allosteric parameters. Finally, we discuss relationships between (thermodynamic constraints on) the allosteric parameters that arise from conformational coupling, and comment on the identifiability of allosteric parameters in receptor oligomers.

## Results

### Receptor oligomers and reduced graph powers

Let *G* = (*V, E*) denote an undirected graph with *v* vertices and *e* edges. Formally, the the vertex set is *V* = {*a*_1_, *a*_2_*,…, a_v_*}, but for readability we will often use the first *v* letters of the alphabet, *V* = {*a, b, c, d*, …}. Each element of the set of edges, *E*, is an unordered pair of vertices. When we say that *G* has the same structure (topology) as a receptor monomer of interest, we mean (*a_i_*, *a_j_*) is an element of *E*(*G*) precisely when there is a reversible transition between states *a_i_* and *a_j_* in the monomer. For a monomer model with *v* states and *e* transitions, *G* will have *v* = *|V|* vertices and *e* = *|E|* edges (using the common notation for the number of elements in a finite set). We assume *G* has no loops or multiple edges, and is connected.

*What graph corresponds to a receptor homomer composed of k identical subunits with topology given by G?* The answer to this question is the *k*th *reduced power of G* (56), denoted by *G*^(*k*)^, which is formally defined as a product graph that is contracted using the symmetries of indistinguishable monomers (see Supporting Material, Sec. S3). For readers with no prior knowledge of product graphs, the state-transition graph of a receptor homo-oligomer can be constructed in 3 steps, as follows.

1. For a receptor model of interest, construct an undirected graph with same topology. For example, an undirected graph *H* = (*V, E*) corresponding to the ternary complex model (Fig. 1) has vertex set *V* = {*a, b, c, d*} and edge wet *E* = (*a, b*), (*a, c*), (*b, d*), (*c, d*) (graph *H* in Fig. 5).
2. Interpreting the vertex labels as variables, write their sum, raise this quantity to the *k*th power, and expand. Each term of the resulting polynomial corresponds to a state of the receptor oligomer. For a dimer composed of *k* = 2 indistinguishable ternary complex monomers, there are 10 distinguishable states

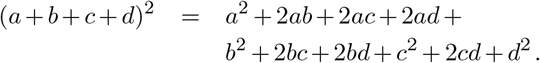 For a ternary complex tetramer, *k* = 4 and (*a* + *b* + *c* + *d*)^4^ = *a*^4^ + *a*^3^*b* + *a*^3^*c* + … + *d*^4^ gives 35 states. In general, the number of states in the receptor oligomer is given by

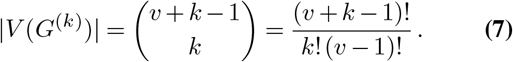 This is the number of ways *k* indistinguishable monomers can each be assigned to one of *v* states.
3. For edges of the receptor oligomer state-transition graph, notice that if (*a_i_*, *a_j_*) is an edge of *G* (an allowed transition in the monomer), there is an edge between two states of *G*^(*k*)^ (a transition in the oligomer) precisely when these states can be written as *a_i_f* = *a_j_f* where *f* (*a*_1_, *a*_2_,…, *a_v_*) is a monomial of degree *k* − 1. The monomial *f* is will be referred to as the *context* of the *a_i_* ⇌ *a_j_* transition, i.e., the unchanged state of *k* − 1 monomers when one monomer changes state from *a_i_* to *a_j_* or vice-versa. Evidently, the number of edges of *G*^(*k*)^ is

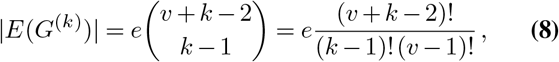

that is, *e* times the number of contexts, which is the number of ways *k* − 1 indistinguishable monomers can each be assigned to one of *v* states. Fig. 5 shows the reduced graph power *H*^(2)^ that gives the topology of the state-transition diagram for a ternary complex homodimer. *H*^(2)^ has 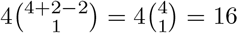 edges. The edge (*ab, bd*) of *H*^(2)^ corresponds to one monomer making an *a* ⇌ *d* transition in the *context* of the other monomer occupying state *b*. Fig. 5 also shows the graph *H*^(4)^ for a receptor oligomer composed of 4 indistinguishable ternary complex monomers. *H*^(4)^ has 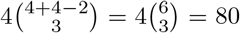 edges. The edge (*b*^2^*cd*, *bc*^2^*d*) of *H*^(4)^ corresponds to one monomer of the receptor 4-mer making an *b* ⇌ *c* transition in the context of *bcd*.

**Fig. 5.**
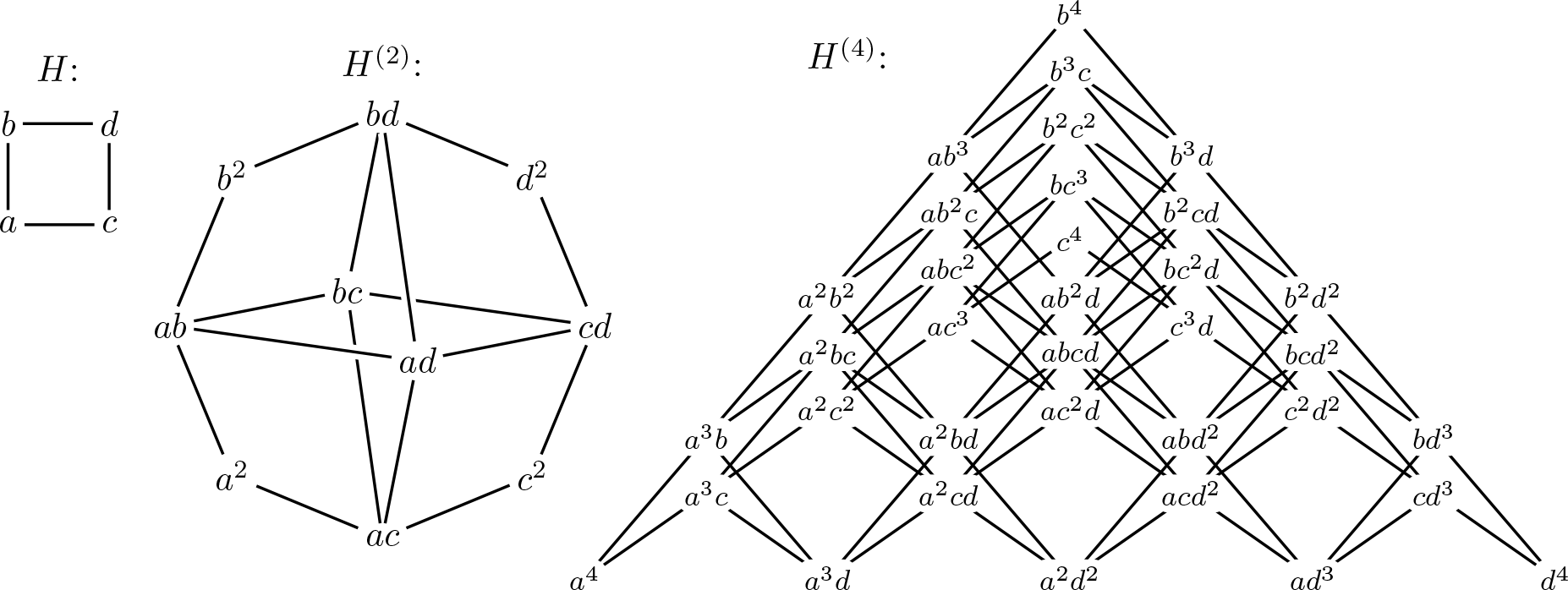
Left: Undirected graph with topology of ternary complex model (Fig. 1). Middle: Topology of a homodimer composed of two *identical and indistinguishable* ternary complex monomers is given by the *reduced graph power H*^(2)^ (see Sec. S3). Right: Topology of a receptor oligomer composed of 4 identical ternary complex monomers is given by the reduced power *H*^(4)^.

### Thermodynamic constraints and the Betti number of *G*^(*k*)^

The number of thermodynamic constraints in a receptor model is given by its *Betti number*, which is the *dimension of the cycle space* of the state-transition graph (for details see Supporting Materials, Sec. S4). Because *G* has no loops or multiple edges, and is connected, its Betti number is given by *β*(*G*) = |*E*(*G*)| − |*V* (*G*)| + 1 = *e* − *v* + 1. The number of free equilibrium parameters in the monomer model is the number of edges less the constraints, which is *e* − *β* = *v* − 1, which is the number of edges in a spanning tree of *G*.

Using Eqs. 7 and 8, the Betti number for the receptor *k*-mer obtained from *G* is *β*(*G*^(*k*)^) = |*E*(*G*)^(*k*)^| − |*V*(*G*)^(*k*)^| +

1. That is, there are

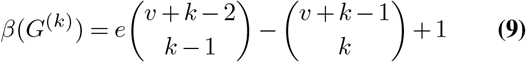

thermodynamic constraints in the oligomer. The ternary complex homodimer has *β*(*H*^(2)^) = 16 − 10 + 1 = 7 thermodynamic constraints and 9 free equilibrium parameters. The 4-mer has *β*(*H*^(4)^) = 80 − 35 + 1 = 46 thermodynamic constraints and 34 parameters (Fig. 5).

### Equilibrium parameters in the monomer model

A general theory of allostery in oligomeric receptors interactions between monomers within the oligomeric receptor begins by introducing a convention for assignment of equilibrium parameters to the edges of *G*, the state-transition graph of the monomer, whose vertex set is *V* (*G*) = {*a*_1_, *a*_2_,…, *a_v_*}. To accomplish this, construct a rooted spanning tree of *G* with root *a*_1_ and indexing that respects a breadth-first traversal (denoted *T*(*G*) or just *T*). Any edge of *T* is uniquely determined by its endpoint *a*_j_ that is furthest from the root. For each 2 ≤ *i* ≤ *v*, let e_*j*_ be the edge of *T* that has endpoints *a_i_* and *a_j_*, with *a_j_* further from the root than *a_i_*. For each edge of *T*, we have 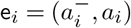, where 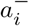 is the predecessor of *a_i_*. For the ternary complex monomer (Fig. 1), an example spanning tree is shown below (left).

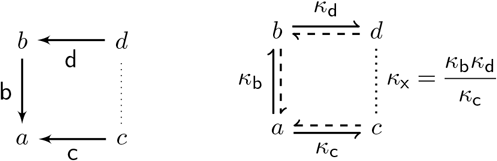

Here the edges are b = (*a, b*), c = (*a, c*), d = (*b, d*) and the predecessors are *b*^−^ = *a*, *c*^−^ = *a*, and *d*^−^ = *b*. The root vertex *a* has no predecessor. For convenience we have chosen *T* so each directed edge e_*i*_ points backwards from product to reactant.

For each edge of *T*, there is a free equilibrium constant that will be denoted by 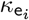 where e_*i*_ is the edge label (above right). For the ternary complex model, the free equilibrium constants are *κ*_b_, *κ*_c_ and *κ*_d_; the constrained equilibrium constant is *κ*_x_ = *κ*_b_*κ*_d_/*κ*_c_. In the notation of Fig. 1, *κ*_b_ = *K_L_*[*L*], *κ*_c_ = *K_G_*[*G*] and *κ*_d_ = *γK_G_*[*G*], and *κ*_x_ = *γK_L_*[*L*].

### Allosteric parameters in oligomeric receptor models

We are now prepared to assign, in a systematic and general fashion, equilibrium parameters to the edges of the receptor oligomer state-transition graph *G*^(*k*)^. Because *T* is a spanning tree of *G*, the reduced power of this spanning tree, denoted *T* ^(*k*)^, spans *G*^(*k*)^ (see Supporting Material, Sec. S6).

As a consequence, a spanning tree of *G*^(*k*)^, denoted Θ(*G*^(*k*)^), can always be constructed using edges that are transition-context pairs involving an edge of *T* (the transition), denoted 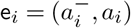, and a monomial *f* (*a*_1_, *a*_2_,…, *a_v_*) of degree *k* − 1 (the context). For now, the equilibrium constants labeling the edges of the spanning tree Θ(*G*^(*k*)^) are formally denoted as 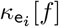 were 2 ≤ *i* ≤ *v*. Using this notation, the equilibrium parameters for the spanning tree Θ(*H*^(2)^) are *κ*_b_[*a*] for the edge (*a*^2^, *ab*), *κ*_b_[*b*] for (*ab, b*^2^), *κ*_d_[*c*] for (*bc, cd*), and so on (Fig. 6A).

**Fig. 6.**
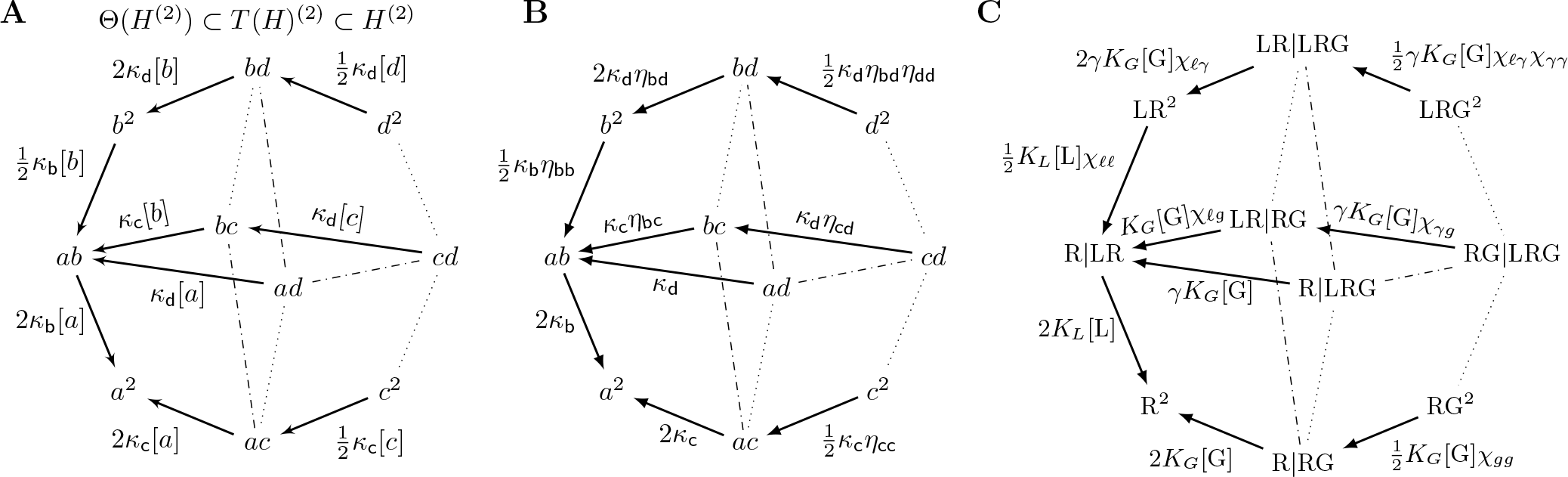
(A) Because *T* is a spanning tree of *H*, the reduced graph product *T* (*H*)^(2)^ spans *H*^(2)^ (but it is not a tree). Θ(*H*^(2)^) is a spanning tree obtained from *T* (*H*)^(2)^ by eliminating three edges (shown dash-dotted). The parameters are transition-context pairs, e.g., *κ*_d_[*b*] denotes the equilibrium constant for reaction d = (*b, d*) in one monomer when the other monomer is in state *b*. (B) The specification of 9 free parameters in the ternary complex homodimer includes 3 equilibrium constants inherited from the monomer (*κ_i_*) and 6 allosteric parameters (*η_i_*). (C) Biophysical notation following Fig. 1 uses the replacements *a → R*, *b →* LR, *c →* RG, *d →* LRG, *η*_bb_ *→ χℓℓ*, *η*_bc_ *→ χℓ_g_*, *η*_bd_ *→ χ*ℓ_*γ*_, *η*_cc_ *→ χ_gg_*, *η*_cd_ *→ χ_gγ_*, *η*_dd_ *→ χ_γγ_* where ℓ, *g* and *γ* stand for R ⇌ LR, R ⇌ RG and LR ⇌ LRG, respectively.

### Enumerating allosteric interactions in receptor dimers

The number of free equilibrium parameters in a receptor oligomer is the number of edges, less the number of thermodynamic constraints, |*E*(*G*^(*k*)^)| − *β*(*G*^(*k*)^) = |*V*(*G*^(*k*)^)| − 1, which is the number of edges in the spanning tree Θ(*G*^(*k*)^). For example, the spanning tree of the homodimer *H*^(2)^, denoted by Θ(*H*^(2)^) and shown in Fig. 6A (solid arrows), is specified by assigning |*E*(Θ(*H*^(2)^))| = 9 parameters. The 4-mer requires |*E*(Θ(*H*^(4)^))| = 34 parameters (Fig. S5, solid arrows).

How should the 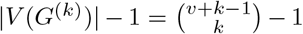 free parameters for a receptor oligomer be specified to illuminate the possible allosteric interactions among monomers? Because the spanning tree *T* (*G*) has *e* = *v* − 1 edges, we may define (*e* + 1)*e*/2 = *v*(*v* − 1)/2 independent 2-way allosteric parameters (the number of ways 2 edges can be chosen from the spanning tree with replacement). For a dimer (*k* = 2), these 2-way parameters are:

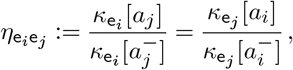

where 2 ≤ *i* ≤ *j* ≤ *v*. For example, the spanning tree *T* (*H*) of the ternary complex monomer has 3 edges (b, c, d). Thus, there are 4 · 3/2 = 6 allosteric parameters for the dimer,

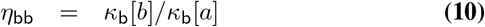

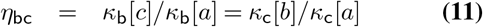

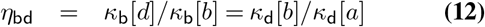

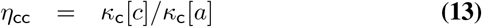

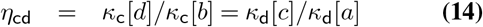

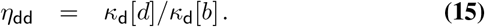

Using the 2-way allosteric parameters defined above, and the equilibrium parameters inherited from the monomer model (*κ*_b_[*a*] = *κ*_b_, *κ*_c_[*a*] = *κ*_c_, *κ*_d_[*a*] = *κ*_d_, because *a* is the root of *T* (*H*)), we are able to specify the equilibrium constants for each edges of Θ(*H*^(2)^) in a manner that illuminates the possibility of conformational coupling. For example, the parameter on edge (*ab, b*^2^) is formally *κ*_b_[*b*], because this edge is a b = (*a, b*) transition in the context of *b*. Using Eq. 10 we have *κ*_b_[*b*] = *κ*_b_[*a*]*η*_bb_ = *κ*_b_*η*_bb_. The edge (*bd, d*^2^) is a d = (*b, d*) transition in the context of *d*. Using both Eqs. 10 and 15, we see that this equilibrium constant is *κ*_d_[*d*] = *κ*_d_[*b*]*η*_dd_ = *κ*_d_*η*_bd_*η*_dd_. Repeating this process for all 10 states yields the specification of allosteric parameters in the ternary complex homodimer shown in Fig. 6B. The corresponding binding curve 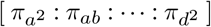 is

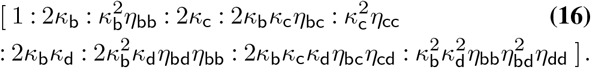

Dividing by (1 + *κ*_b_ + *κ*_c_ + *κ* _b_*κ*_d_)^2^ gives

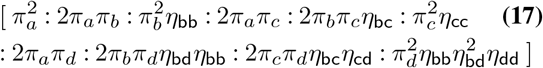

where [*π_a_*: *π_b_*: *π_c_*: *π_d_*] = [1: *κ*_b_: *κ*_c_: *κ*_b_*κ*_d_].

As promised, without assuming independence of receptor monomers, we have expressed the equilibrium occupancy measure of the ternary complex homodimer in terms of the properties of an isolated monomer (*π_a_*, *π_b_*, etc.) and allosteric parameters (*η*_bb_, *η*_bc_, etc.). Fig. 6C shows the distinguished spanning tree of the homodimer in the original biophysical notation (Fig. 1) and Sec. S2 in the Supporting Material presents the biophysical version of Eq. 16.

### Multiway allosteric interactions

For receptor oligomers with *k >* 2, the 2-way parameters are

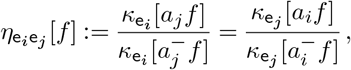

where 2 ≤ *i* ≤ *j* ≤ *v* and *f*(*a*_1_, *a*_2_,…, *a_v_*) is a monomial of degree *k* − 2. Furthermore, when when *k* > 2, the situation is complicated by the possibility of multiway allosteric interactions. For example, the 3-way allosteric parameters are

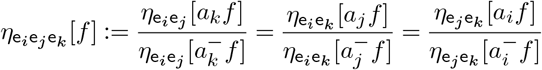

where 2 ≤*i* ≤ *j* ≤ *k* ≤ *v* and *f* (*a*_1_, *a*_2_,…, *a_v_*) is a monomial of degree *k* − 3(when *k* = 3 the *f* is dropped). The equalities are shown by expanding the definition,

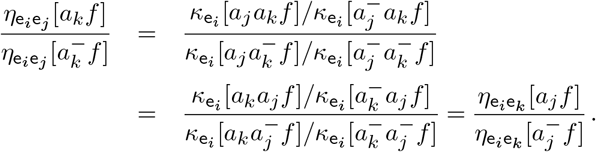

In general, *n*-way allosteric parameters are defined as

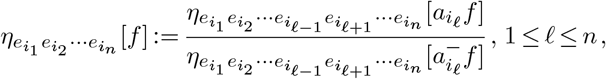

where 2 ≤ *i*_1_ ≤ *i*_2_ ≤ · · · ≤ *i_n_* ≤ *v*. For a monomer with spanning tree of *e* = *v* − 1 edges, a receptor composed of *k* monomers has 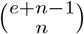 *n*-way allosteric parameters for 2 ≤ *n* ≤ *k*, which is the number of ways that *n* of the *e* edges can be chosen with replacement. For example, the spanning tree *Theta*(*H*^(4)^) of the ternary complex 4-mer has 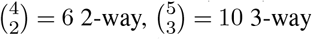 and 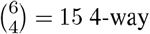 parameters. Some of these are

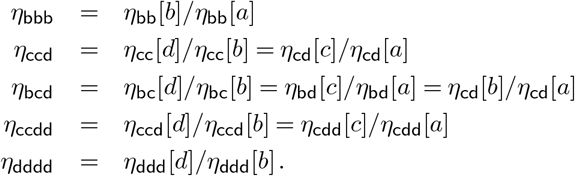

### Token method for allosteric parameters

Fortunately, the allosteric factors involving various 2-way through *n*-way interactions may be enumerated using a natural ‘token’ representation of receptor oligomer states, as follows.

To begin, draw the tree *T* that spans the state-transition graph *G* of the monomer (discussed above). For any given state of the oligomer, put (indistinguishable) tokens in the positions associated with the monomer states. For example, the token graphs associated to states *bd* and *d*^2^ in the ternary complex dimer are,

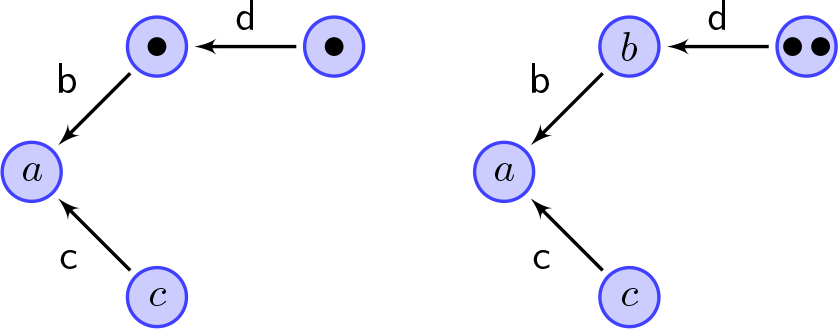

To calculate allosteric factor for state *bd*, we consider the path of each token to the root (the vertex *a*). These paths yield b for the first token, and b+d for the second token. Because the product is b(b+ d) = b^2^ + bd, the allosteric factor in the term] *π_b_π_d_* in Eq. 17 is *η*_bb_*η*_bd_. For state *d*^2^, the path to root for both tokens is *b* + *d* and the product is 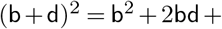 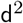; thus, the allosteric factor for 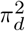 is 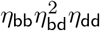. Table S1 shows the complete list of allosteric factors for the ternary complex dimer.

For a receptor *k*-mer, we may assume *p*_1_ ≤ *p*_2_ ≤ … ≤ *p*_k_ where *p*_ℓ_ is the place of the ℓth token. Recall that 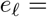 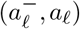 and define *h*(*a*_ℓ_) recursively,

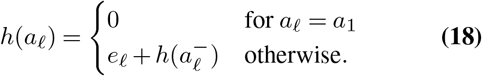

The *n*-way interactions are enumerated by the elementary symmetric polynomials in *h*_1_, *h*_2_,…, *h_k_*, namely,

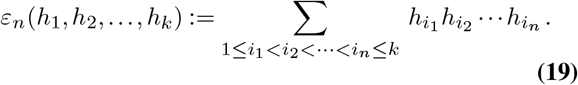

For example, the token graphs associated to states *bcd*^2^ and *ac*^2^*d* in the ternary complex 4-mer are,

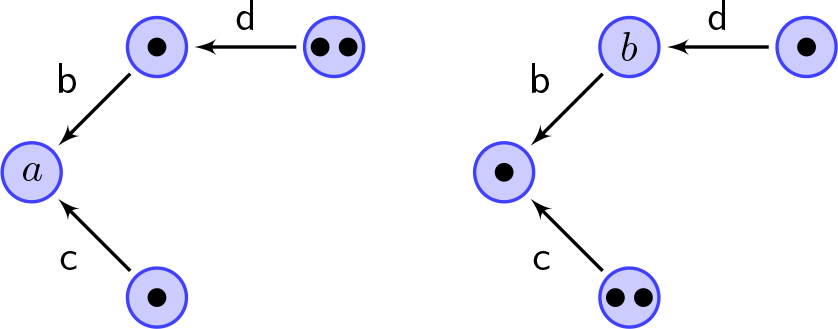

The 2-, 3- and 4-way interactions (Eqs. 18 and 19) are

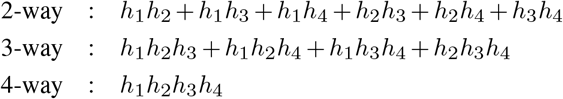

where

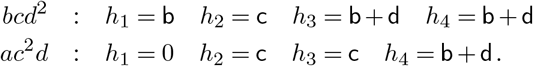

For state *bcd*^2^,

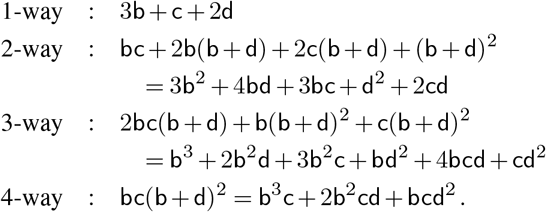

Thus, the 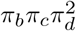 term, which has combinatorial coefficient (1, 1, 2)! = 4!/(0!1!1!2!) = 12, has allosteric factors

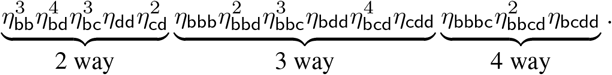

For state *ac*^2^*d*, a similar calculation gives

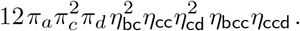

Sec. S7 confirms that the token method for enumerating allosteric factors in a receptor oligomer always yields the required 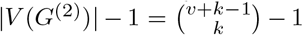 parameters. Because the spanning tree *T*(*G*) used to define the |*V*(*G*)| − 1 unconstrained equilibrium constants in an isolated monomer is not unique. Comparing results for different choices of *T* reveals identities that relate allosteric parameters. In the ternary complex dimer (Fig. 6), e.g., *η*_bb_*η*_bd_ = *η*_bc_*η*_bx_, where *η*_bxb_:= *κ*_b_[*d*]/*κ*_b_[*c*] = *κ*_x_[*b*]/*κ*_x_[*a*] and *κ*_x_[*a*] = *κ*_b_*κ*_d_/*κ*_c_ (see Fig. S4).

## Discussion

The theoretical framework for understanding allostery in receptor oligomers presented here represents an intriguing and novel combination of graph theory and quantitative receptor pharmacology. We began by establishing that the structure of state-transition diagram of a receptor *k*-mer, for any given monomer topology *G*, is the reduced graph power *G*^(*k*)^. We used a minimal cycle basis construction for reduced graph powers to identify thermodynamic constraints in receptor oligomers *without having to construct G*^(*k*)^. We showed how allosteric interactions in receptor oligomers may be systematically enumerated. Finally, we show how the occupancy measure of a receptor oligomer may be expressed in terms of the parameters for an isolated monomer and these identified allosteric parameters (Eq. 17).

The concepts and notation introduced here amount to a *theoretical framework for allostery in oligomeric receptors composed of any number of identical monomers*. For clarity we have used the (perhaps over-simple) ternary complex model dimer and tetramer as running examples, but the approach is completely general. See Fig. 8 and Sec. S8 for discussion of cubical ternary complex dimers (49). However, the approach is completely general. For any given spanning tree *T* (*G*) of a monomer state transition diagram *G* that is of interest, the allosteric parameters can be enumerated by performing the symbolic calculations of Eqs. 18 and 19 in a computer algebra system (see Sec. S9 in the Supporting Material).

We hope this theoretical framework for receptor homomer allostery will be valuable to investigators interested in pharmacological alteration of GPCR activity by allosteric modulators, whose action is modeled as a modification of equilibrium constants of one or more receptor monomers. Using a similar approach, an understanding of allosteric interactions in hetero-oligomers is within reach, which could lay a foundation for theoretical analysis of receptor crosstalk, wherein one protomer binds to an agonist, whilst the other unit activates the G protein (57, 58).

Using this framework, it may be possible to address, in a general fashion, the identifiability (or not) of allosteric parameters (*η_i_*) that emerge in receptor oligomer models. The outcome of such studies would presumably depend on whether the equilibrium parameters inherited from the monomer (*κ_i_*) have been experimentally validated and, consequently, are fixed during the process of fitting allosteric parameters to experimental data sets. The limitations of this theoretical framework for allostery in receptor oligomers need not be enumerated, as these are inherited from the limitations of the general and accepted practice of receptor occupancy modeling.

**Fig. 7.**
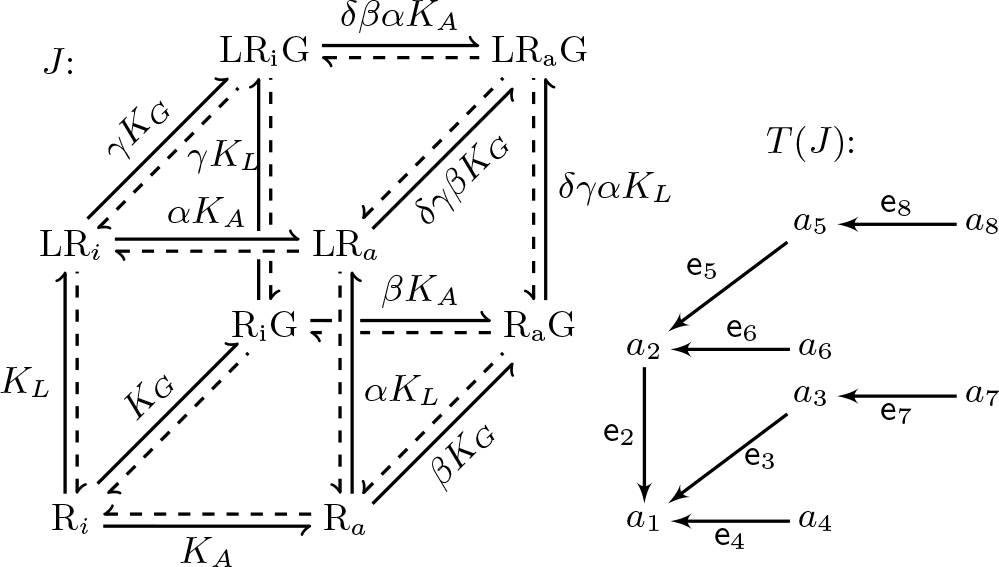
Left: Cubical ternary complex model of a G protein coupled receptor (49). There are 7 equilibrium parameters: two associations constants (*K_L_*, *K_G_*) for the binding of ligand and G protein, one equilibrium constant (*K_A_*) for receptor activation, and four allosteric parameters (*α*, *β*, *γ*, *δ*). Right: The spanning tree *T* (*J*) respecting breadth first traversal is the starting point for constructing a distinguished spanning tree Θ(*J*^(2)^) of the cubical ternary complex dimer (Fig. 8)

**Fig. 8.**
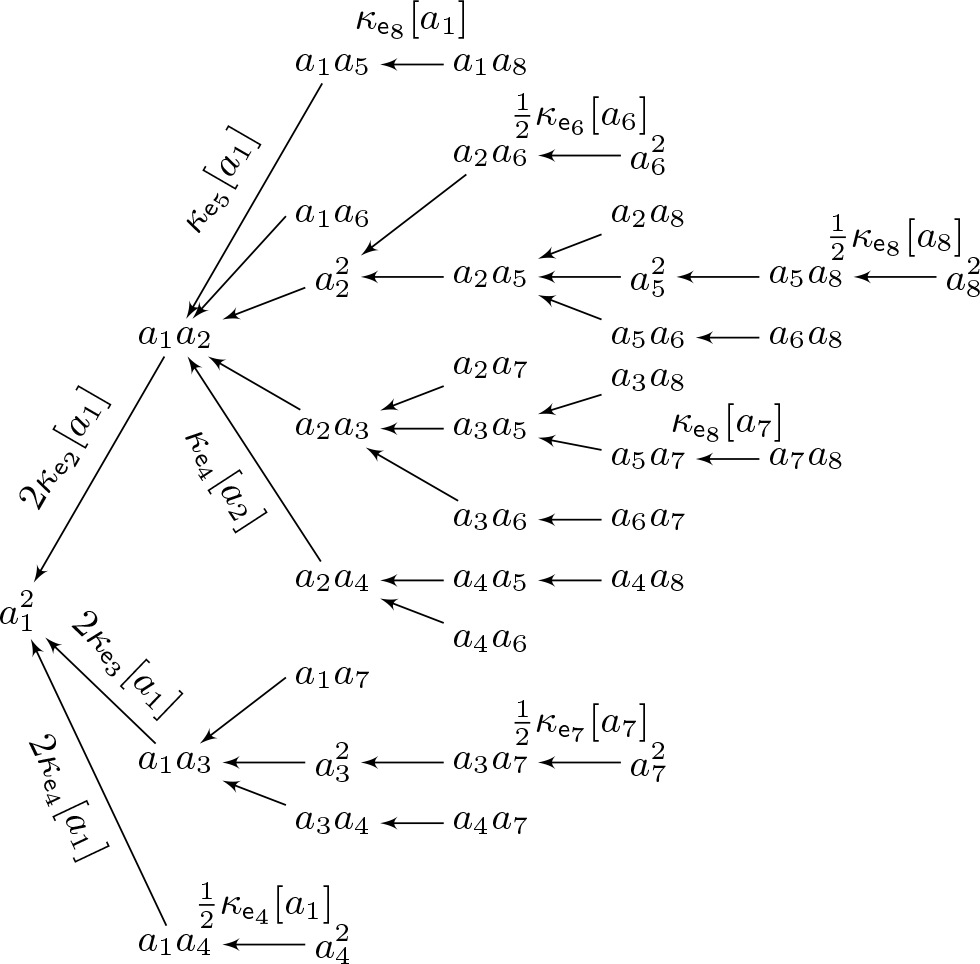
Distinguished spanning tree Θ(*J*^(2)^) for the cubical ternary complex homodimer (Fig. 7). Fig. S6 in the Supporting Materials shows state-transition diagram *J*^(2)^. Fig. 7 shows |*V* (*J*)| = 8 so 7 parameters are inherited from the monomer (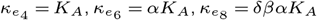, etc.). The dimer has |*V* (*J*^(2)^)| = 36 vertices; thus, there are 35 parameters, 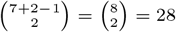 of which are 2-way allosteric parameters 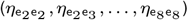. Relationship between formal and specified equilibrium parameters include 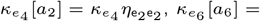 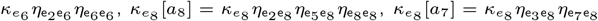, among others (see Sec. S8 in Supporting Material)

## Acknowledgements

GDCS is grateful for numerous stimulating conversations with Ron Smith, Dan Borrus and Wenchong He. The MCB construction for reduced graph powers (Eqs. S27–S29) was found in collaboration with Richard H. Hammack and its description in the Supplementary Notes is adapted from (56).

## Supplementary Note S1: Example binding curves for the ternary complex model

For any [L] and [G], the equilibrium fraction of receptors in each of the four states of the ternary complex model (Fig. 1) can be found by expressing each receptor state concentration in terms of [R],

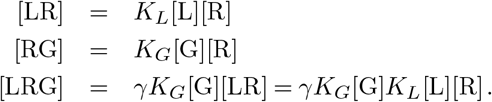

Solving these equations simultneously with the equation for the conserved total receptor concentration, namely,

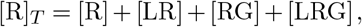

gives the fraction of receptors in each state,

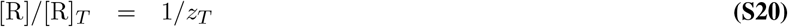

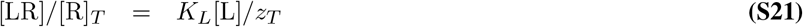

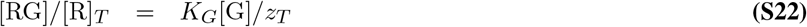

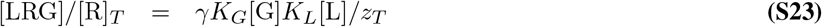

where *z_T_* = 1 + *K_L_*[L] + *K_G_*[G] + *γK_G_*[G]*K_L_*[L].

**Fig. S1.**
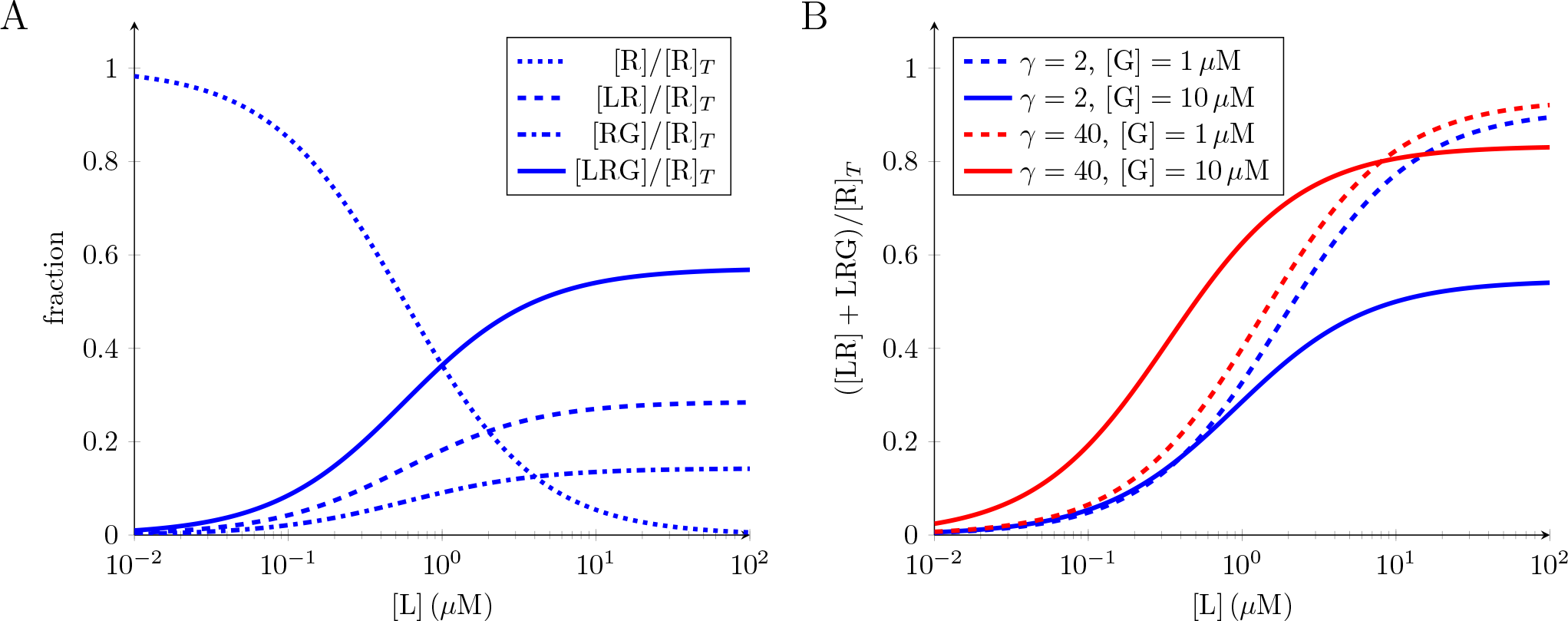
Binding curves for ternary complex model (Eqs. S20–??). Parameters: *K_L_* = 0.5*μ*M^−1^, *K*_*G*_ = 0.1*μ*M^−1^, and in (A) [G] = 5*μ*M and *γ* = 40

**Fig. S2.**
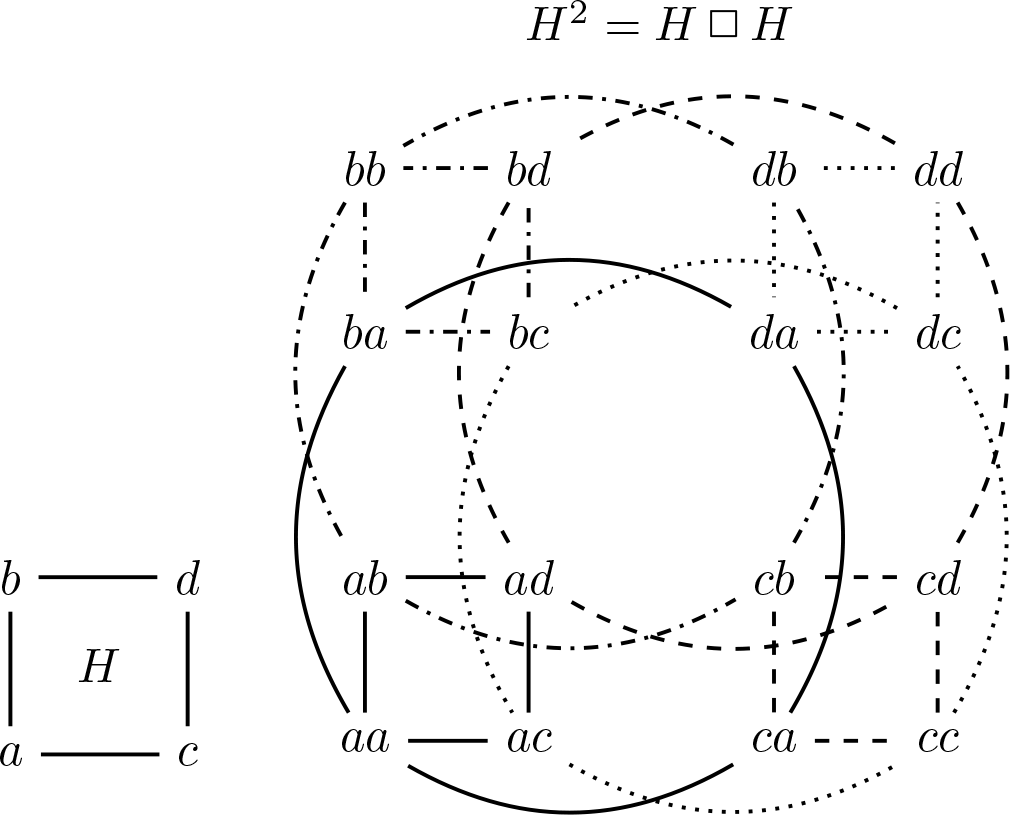
Topology of a receptor dimer composed of two distinguishable ternary complex monomers (*H*, see Fig. 1) is given by the Cartesian graph power *H*^2^ = *H*◻*H*

## Supplementary Note S2: Ternary Complex Dime

In the original biophysical notation, the fraction of dimeric receptors in each state can be read off Fig. 6C, as follows.

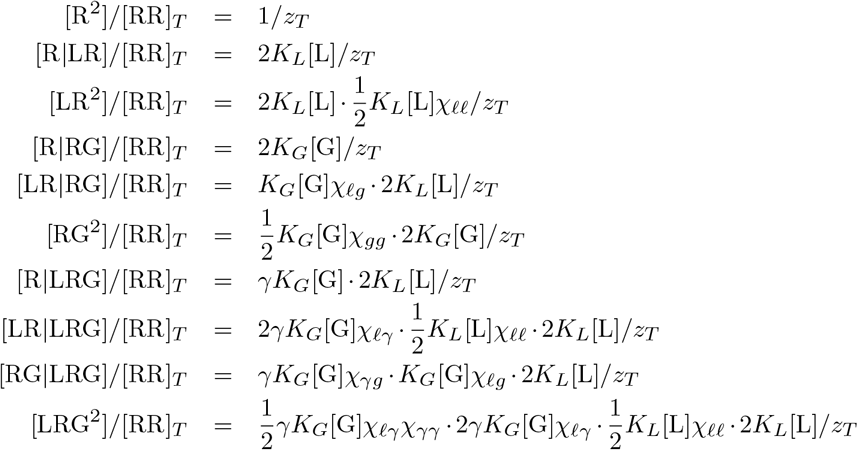

where

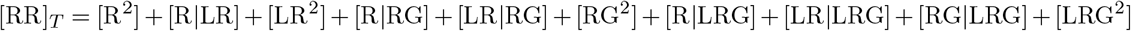

and *z_T_* is the sum of the right sides of the above equations. Equivalently,

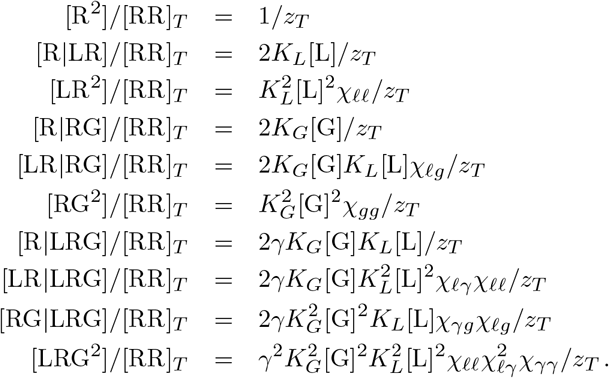

**Fig. S3.**
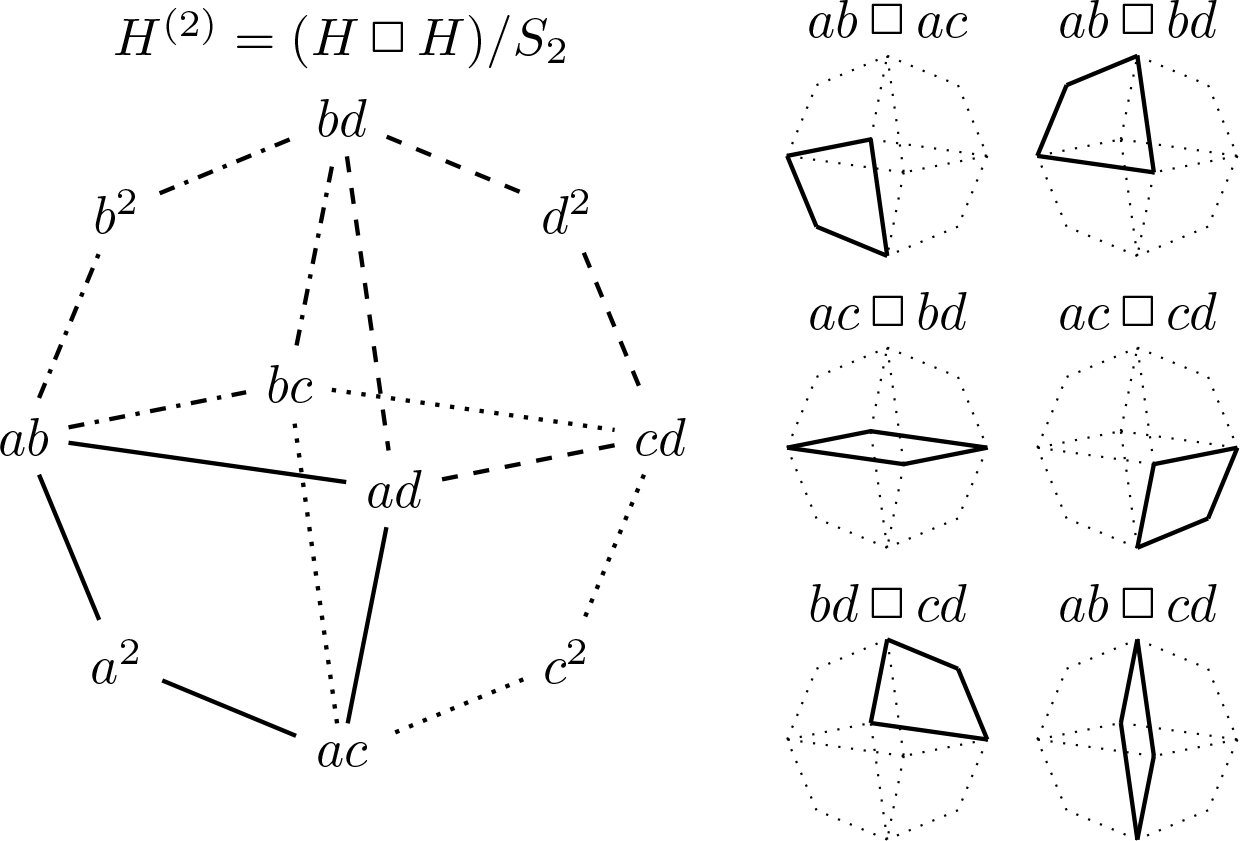
Topology of a homodimer composed of two *identical and indistinguishable* ternary complex monomers (*H*, see Fig. 1) is given by the *reduced graph power H*^(2)^ (bottom). For each vertex *x* of *H*, the vertices {*xv* | *v* ∈ *V* (*H*)} ⊆ *V* (*H*^(2)^) induce a subgraph *Hx* ≅ *H* of *H*^(2)^. The subgraph *Ha* is shown solid in *H*^(2)^. Right: 6 of 10 4-cycles in *H*^(2)^ are Cartesian squares

## Supplementary Note S3: Receptor oligomers and reduced graph powers

Let the undirected graph *G* = (*V* (*G*), *E*(*G*)) represent a receptor subunit model with *v* = |*V*| (*G*) states and *e* = |*E*(*G*)| reversible transitions between these states. We assume *G* has no loops or multiple edges, and is connected. *What graph corresponds to a receptor homomer composed of k identical subunits with topology given by G?* The answer to this question is the *k*th *reduced power of G*, denoted by *G*^(*k*)^, which is defined as a contraction of product graphs. We briefly review this construction, following prior work (56).

Recall (59) that the *Cartesian product* of graphs *G* and *H* is the graph *G*◻*H* whose vertex set is the Cartesian product *V* (*G*) × *V* (*H*), and whose edges are

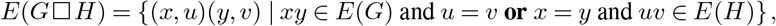

where *xy* is an abbreviation for a vertex (*x, y*) of *G*◻*H* and, similarly, *uv* = (*u, v*) ∈ *V* (*G*◻*H*). The *k*th *Cartesian power* of *G* is the *k*-fold product *G^k^* = *G*◻*G*◻ … ◻ *G*. The *reduced k*th power is the quotient *G^k^/S_k_* of *G*^*k*^ by the action of the symmetric group *S_k_*, which acts on *G^k^* by permuting the factors.

In general, say *G* has vertex set {*a*_1_, *a*_2_,…, *a_v_*}. Denote by *M_k_* (*G*) the set of monic monomials of degree *k*, with indeterminates *V* (*G*), and *M*_0_(*G*) = {1}. Then *G*^(*k*)^ is the graph whose

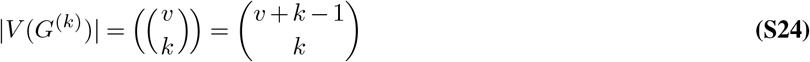

vertices are the monomials 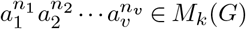. For edges, if *a_i_a_j_* is an edge of *G*, and *f* (*a*_1_, *a*_2_,… *a_v_*) ∈ *M*_*k*−1_(*G*), then *a_i_f* (*a*_1_, *a*_2_,…, *a_v_*) is adjacent to *a_j_f* (*a*_1_, *a*_2_,…, *a_v_*). For each edge *a_i_a_j_* of *G* and each monomial *f* ∈ *M*_*k*−1_(*G*), there is an edge of *G*^(*k*)^ from *a_i_f* to *a_j_f*; thus, the number of edges of *G*^(*k*)^ is

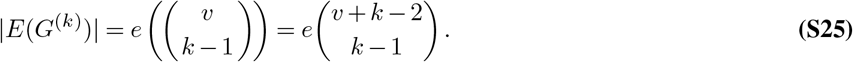

As an example, consider the graph *H* shown in Fig. S2, which has the topology of the ternary complex model shown in Fig. 1. Here *V* (*H*) = {*a, b, c, d*} and *E*(*H*) = {{*a, b*}, {*a, c*}, {*b, d*}, {*c, d*}}. The graph power *H*^2^ has *v*^2^ = 16 vertices and 2*ev* = 32 edges. This should be compared to Fig. 5, which shows the *reduced* graph power *H*^(2)^ that corresponds to the state-transition diagram of a ternary complex receptor dimer composed of *identical and indistinguishable* monomers. *H*^(2)^ has 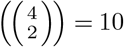 vertices and 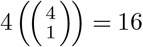 edges. Fig. 5 shows the reduced graph product *H*^(4)^ that corresponds a receptor oligomer composed of 4 ternary complex monomers. *H*^(4)^ has 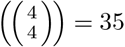 vertices and 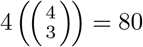 edges. The vertices enumerated by expanding the multinomial (*a* + *b* + *c* + *d*)^4^ and dropping coefficients, because *V*(*H*^(4)^) = *M*_4_(*H*).

## Supplementary Note S4: Cycle space and thermodynamic constraints in receptor oligomer models

The number of thermodynamic constraints in a receptor model is given by *dimension of the cycle space* of the state-transition graph. Following (56), we briefly review this concept.

For a graph *G*, its *edge space* ɛ(*G*) is the power set of *E*(*G*) viewed as a vector space over the two-element field 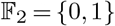, where the zero vector is 0 = ∅ and addition is symmetric difference. Any vector *X* ∈ ɛ(*G*) is viewed as the subgraph of *G* induced on *X*, so ɛ(*G*) is the set of all subgraphs of *G* without isolated vertices. Thus *E*(*G*) is a basis for ɛ(*G*), and dim(ɛ(*G*)) = |*E*(*G*)|. For a graph *G*, its *cycle space* 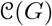 is the set of all subgraphs in ɛ(*G*) whose vertices all have even degree (that is, the Eulerian subgraphs). Because every such subgraph can be decomposed into edge-disjoint cycles, we see that 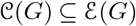 is spanned by the cycles in *G*. The dimension of 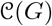 is called the *Betti number* of *G*, and is denoted *β*(*G*). If *G* is connected, then *β*(*G*) = |*E*(*G*)| − |*V* (*G*)| + 1 (e.g., *β*(*H*) = 1 in Fig. S2). A basis for 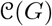 is called a *cycle basis* for *G*. The *length* of a cycle basis is the sum of the lengths of its cycles. A cycle basis with the least possible length is a *minimum cycle basis* (MCB). See Chapter 29 of (59) for further review.

For a receptor oligomer, the number of thermodynamic constraints is the dimension of the cycle space of the reduced graph product, 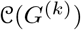. Using Eqs. 7 and 8, this is given by the Betti number

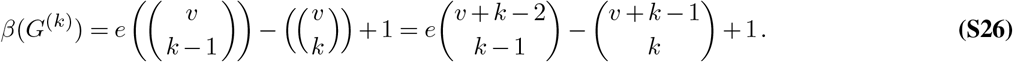

## Supplementary Note S5: Minimal cycle basis of reduced graph powers

Enumerating the *β*(*G*^(*k*)^) thermodynamic constraints of a receptor oligomer model is achieved by decomposing cycle space of a reduced power *G*^(*k*)^ into the direct sum of two particularly simple subspaces. Following (56), we briefly describe this process.

1. First, notice that if *f* is a fixed monomial in *M*_*k*−1_(*G*), then *x ↦ xf* is an embedding *G → G*^(*k*)^. Call the image of this map *G f*. Note that *G f* is an induced subgraph of *G*^(*k*)^ and is isomorphic to *G*.
2. Next, we define a special type of cycle in a reduced power. Given distinct edges *wx* and *yz* of *G* and any *f* ∈ *M*_*k*−2_(*G*), there is a square in *G*^(*k*)^ with vertices *wyf, xyf, xzf, wzf*. Let us call such a square a *Cartesian square*, and denote it as (*wx*◻*yz*)*f*. Note that a subgraph *Gf* of *G*^(*k*)^ may have squares (4-cycles), but these are not *Cartesian squares*, because they do not have the form specified above. In Fig. S3, *Ha* = *a*^2^ + *ab* + *ac* + *ad ⊂ H*^(2)^ is *not* a Cartesian square, whereas the square *ab*◻*bd* = *ab* + *b*^2^ + *bd* + *ad is* a Cartesian square.
3. Define the *square space* 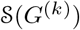 to be the subspace of 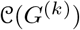 that is spanned by the Cartesian squares. Any pair of distinct edges *wx* and *yz* of *G* corresponds to a Cartesian square (*wx*◻*yz*)*f*, where *f* ∈ *M*_*k*−2_(*G*), so there are 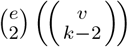 such squares; however, this set of squares may not be independent. Prior work (56) gives a construction of a square basis, i.e., a maximum independent set 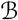 of Cartesian squares in 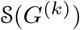 (see below).
4. For fixed *f* ∈ *M*_*k*−1_(*G*), the cycle space of reduced *k*th power of *G* is the direct sum

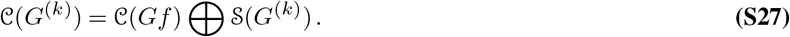

Thus, a basis for 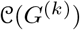 may be constructed from independent cycles in 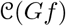 and 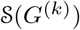, as follows. Take a cycle basis 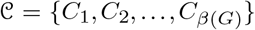 for *G*, and let 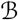 be a square basis for 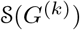. Fix a monomial *f* ∈ *M*_*k*−1_(*G*) and put 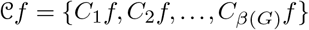. Then 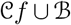 is a cycle basis for *G*^(*k*)^. If 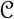 is a minimal cycle basis (MCB) for *G*, and *G* has no triangles, then this basis is an MCB for *G*^(*k*)^.

The *square basis* 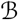 may be constructed as follows. Put *V* (*G*) = *a*_1_, *a*_2_,…, *a_v_*. Let *T* be a rooted spanning tree of *G* with root *a*_1_ with indexing that respects a breadth-first traversal. Any edge of *T* is thus uniquely determined by its endpoint *a_j_* that is furthest from the root. For each 2 ≤ *i* ≤ *v*, let e_*j*_ be the edge of *T* that has endpoints *a_i_* and *a_j_*, with *a_j_* further from the root than *a_i_*. Let *M*_*k*−2_(*a*_1_, *a*_2_,… *a_j_*) denote the monic monomials of degree *k* − 2 in indeterminates *a*_1_, *a*_2_,…, *a_j_*, with 1 ≤ *j* ≤ *v*.

Define the following sets of Cartesian squares in *G*^(*k*)^:

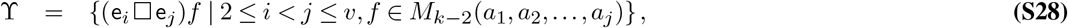

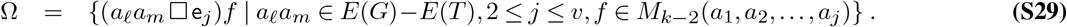

The set 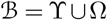 is a basis for the square space 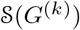. Note: in the above expressions, when *j* < *v*, *M*_*k*−2_(*a*_1_, *a*_2_,…, *a_j_*) ⊂ *M*_*k*−2_(*a*_1_, *a*_2_,…, *a_v_*), and the monomials *f* in Eqs. S28 and S29 do not involve the variables *a*_*j*+1_, *a*_*j*+2_,…, *a_v_*.

## Supplementary Note S6: Identifying thermodynamic constraints

The minimal cycle basis construction for the reduced graph power *G*^(*k*)^ presented above suggests a systematic method for enumerating *β*(*G*^(*k*)^) edges of a receptor oligomer state-transition graph that correspond to thermodynamic constraints (Eq. S26).

The remaining |*V* (*G*^(*k*)^)| edges will be a distinguished spanning tree of *G*^(*k*)^, denoted by Θ(*G*^(*k*)^), to which we will assign equilibrium parameters.

1. Choose a rooted spanning tree of the monomer (*T ⊂ G*) with indexing that respects a breadth-first traversal (as in point (3) of the previous section). Because *T* has no cycles, *β*(*T*) = 0 and 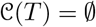. Note that the edges of the spanning tree *T* are directed reactants ← products (arrow oriented as the reverse reaction). Let 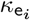 denote the equilibrium parameter associated with edge e_*i*_ ∈ *E*(*T*).
2. Construct the reduced graph power *T*^(*k*)^, a reduced graph power with the same vertex set as *G*^(*k*)^, namely, *M*_*k*_(*a*_1_, *a*_2_,…, *a_v_*). The edges of *T*^(*k*)^ inherit direction from *T*, because edges of *T*^(*k*)^ take the form e_*i*_*f* where *f* ∈ *M*_*k*–1_. Note that *T*^(*k*)^ spans *G*^(*k*)^, and *E*(*T*^(*k*)^) = *E*(*G*^(*k*)^) − Ω with Ω as in Eq. S29. Thus, the cycle space of *T*^(*k*)^ is its square space, 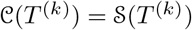 in Eq. S27, and a basis for 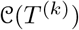 may be constructed from the set ϒ (Eq. S28) of independent cycles in 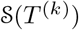.
3. The distinguished spanning tree Θ(*G*^(*k*)^) we seek has edges

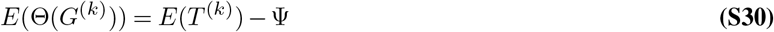

where Ψ is a set of distinct edges, one from each square in ϒ, as follows. Write 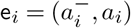 where 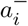 is the predecessor of *a_i_* in *T* (and similarly for e_*j*_). Each square (e_*i*_◻e_*j*_)*f* ∈ ϒ is composed of the four edges (Eq. S28),

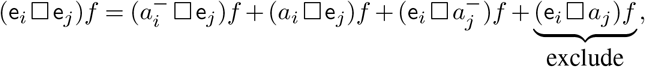

where 2 ≤ *i* < *j* ≤ *v* and *f* ∈ *M*_*k*−2_(*a*_1_, *a*_2_,…, *a_j_*). These edges are directed as e_*i*_ and e_*j*_,

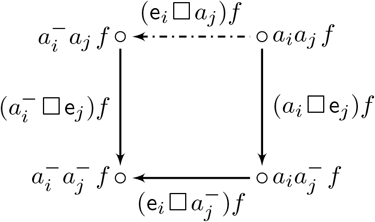

We include in Φ—and, thus, exclude from Θ(*G*^(*k*)^)—the edge (e_*i*_◻*a_j_*)*f*, shown dashdotted above, that is the transition-context pair with context *a_j_* that is last in dictionary order (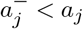 and 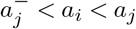). Thus, the set Ψ in Eq. S30 is

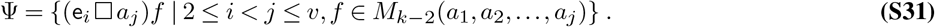

These edges are distinct, and |Ψ| = β(*T*^(*k*)^).

In this way—following steps (1)–(3) above—we construct Θ(*T*^(*k*)^), a subgraph of *T* ^(*k*)^ that is a distinguished spanning tree of *G*^(*k*)^. Two examples follow.

### A. Ternary complex dimer

For *H*^(2)^, shown in Fig. 5, *k* = 2 and *M*_*k*−2_ = {1}. Eq. S28 and the rooted spanning tree *T ⊂ H* shown in Fig. 6 yields the following basis of the square space 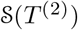,

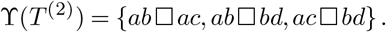

The 12 edges of these 3 Cartesian squares are

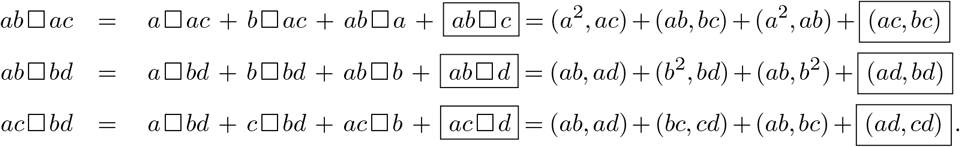

The boxed edges are Ψ = (*ac, bc*), (*ad, bd*), (*ad, cd*), i.e., the edges of *T*^(2)^ that are not included used in the spanning tree Θ(*T*^(2)^). These excluded edges are shown dashdotted in Fig. 6. The spanning tree Θ(*G*^(2)^) has edges *E*(*T*^(*k*)^) − Ψ (Eq. S30).

Fig. S4 provides insight into how the specification of allosteric parameters in the ternary complex homodimer depends (superficially) on the chosen rooted tree *T* (*G*) that spans the monomer state-transition diagram *G*.

**Fig. S4.**
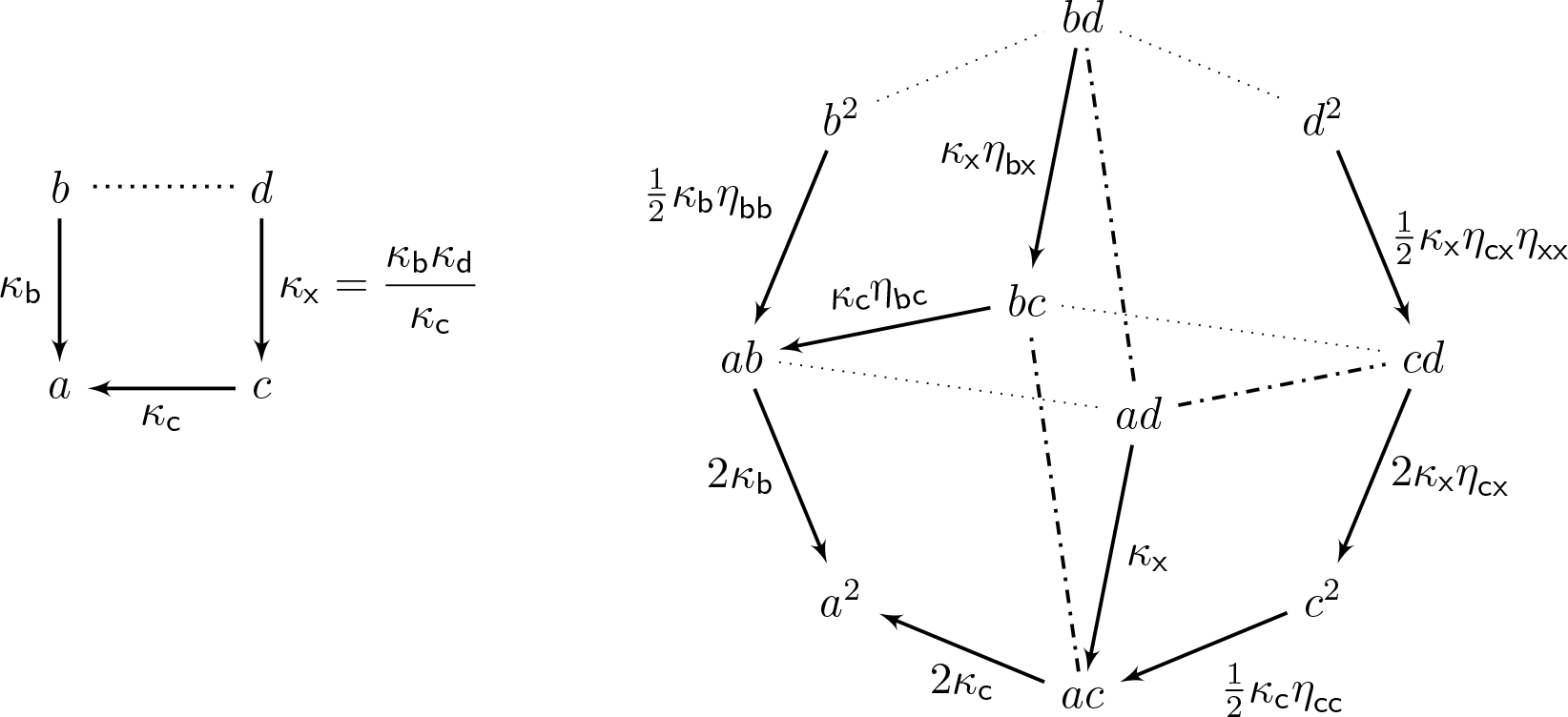
Specification of allosteric parameters in the ternary complex homodimer depends (superficially) on the chosen rooted tree *T* (*G*) that spans the monomer state-transition diagram *G*. Comparing the paths *bd* → *bc* → *ab* and *bd* → *b*^2^ → *ab* using the specification shown here and Fig. 6B reveals that 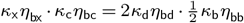. Because *κ*_x_ *κ*_c_ = *κ*_d_ *κ*_b_, we discover that *η*_bx_ *η*_bc_ = *η*_bb_ *η*_bd_. This is true by definition, because *η*_bx_ = *κ*_b_[*d*]/*κ*_b_[*c*], *η*_bc_ = *κ*_b_[*c*]/*κ*_b_[*a*], *η*_bb_ = *κ*_b_[*b*]/*κ*_b_[*a*], *η*_bd_ = *κ*_b_[*d*]/*κ*_b_[*b*]. Another example: comparing the paths *cd → c*^2^ *→ ac* and *cd → ad → ac* reveals that *η*_cx_ *η*_cc_ = *η*_bc_ *η*_cd_

### B. Ternary complex 4-mer

For *H*^(4)^, shown in Fig. 5, the Betti number is 46. Fig. S5 shows the spanning tree *T ⊂ H* and the reduced graph product *T*^(4)^ ⊂ *H*^(4)^, which spans *H*^(4)^ and has Betti number 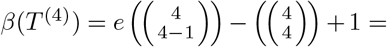 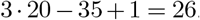. The basis ϒ of the square space (and cycle space) of *T*^(4)^ is the following set of Cartesian squares,

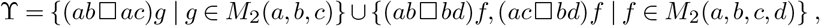

where *M*_2_(*a, b, c*) = {*a*^2^, *ab, b*^2^, *ac, bc, c*^2^} and *M*_2_(*a, b, c, d*) = *M*_2_(*a, b, c*) ∪ {*ad, bd, cd, d*^2^}. Using *ab*◻*c* = (*ac, bc*), *ab*◻*d* = (*ad, bd*), and *ac*◻*d* = (*ad, cd*), the 6 + 2 *·* 10 = 26 element set Ψ, shown dashdotted in Fig. S5, is

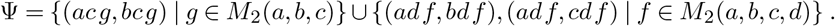

The graph Θ(*G*^(*k*)^), the subgraph of *T* ^(*k*)^ that is the distinguished spanning tree of *G*^(*k*)^, has edges *E*(*T*^(*k*)^) − Ψ and is shown solid in Fig. S5.

## Supplementary Note S7: Counting allosteric parameters

The token method for enumerating each allosteric factor in a receptor oligomer model always yields the required 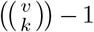 parameters. To see this, note that the number of *n*-way allosteric interactions involving the *v* − 1 = *e − β*(*G*) edges of the spanning tree *T* ⊂ *G* is 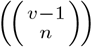 When *n* = 1 this is 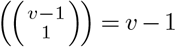 (the number of free parameters in a monomer). When these are supplemented with the 2-way through *k*-way interactions, the correct number of parameters is obtained,

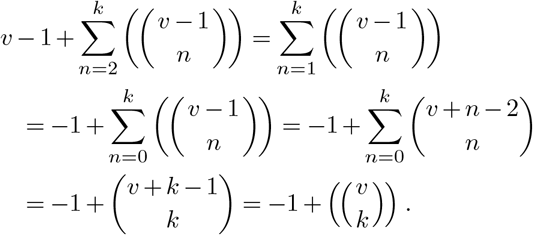

**Table S1.** Worksheet for the token method for enumerating allosteric parameters in the case of the ternary complex dimer (compare Fig. 6B and Eq. 17).

| state | token 1 | token 2 | sum | 1-way | product | 2-way |
| --- | --- | --- | --- | --- | --- | --- |
| $a^2$ | 0 | 0 | 0 | 0 | 0 | - |
| $ab$ | 0 | b | b | $\kappa_b$ | 0 | - |
| $b^2$ | b | b | 2b | $\kappa_b^2$ | $b^2$ | $\eta_{bb}$ |
| $ac$ | 0 | b + c | b + c | $\kappa_b \kappa_c$ | 0 | - |
| $bc$ | b | b + c | 2b + c | $\kappa_b^2 \kappa_c$ | $b^2 + bc$ | $\eta_{bb} \eta_{bc}$ |
| $c^2$ | b + c | b + c | 2b + 2c | $\kappa_b^2 \kappa_c^2$ | $b^2 + 2bc + c^2$ | $\eta_{bb} \eta_{bc}^2 \eta_{cc}$ |
| $ad$ | 0 | d | d | $\kappa_d$ | 0 | - |
| $bd$ | b | d | b + d | $\kappa_b \kappa_d$ | bd | $\eta_{bd}$ |
| $cd$ | b + c | d | b + c + d | $\kappa_b \kappa_c \kappa_d$ | bd + cd | $\eta_{bd} \eta_{cd}$ |
| $d^2$ | d | d | 2d | $\kappa_d^2$ | $d^2$ | $\eta_{dd}$ |

**Fig. S5.**
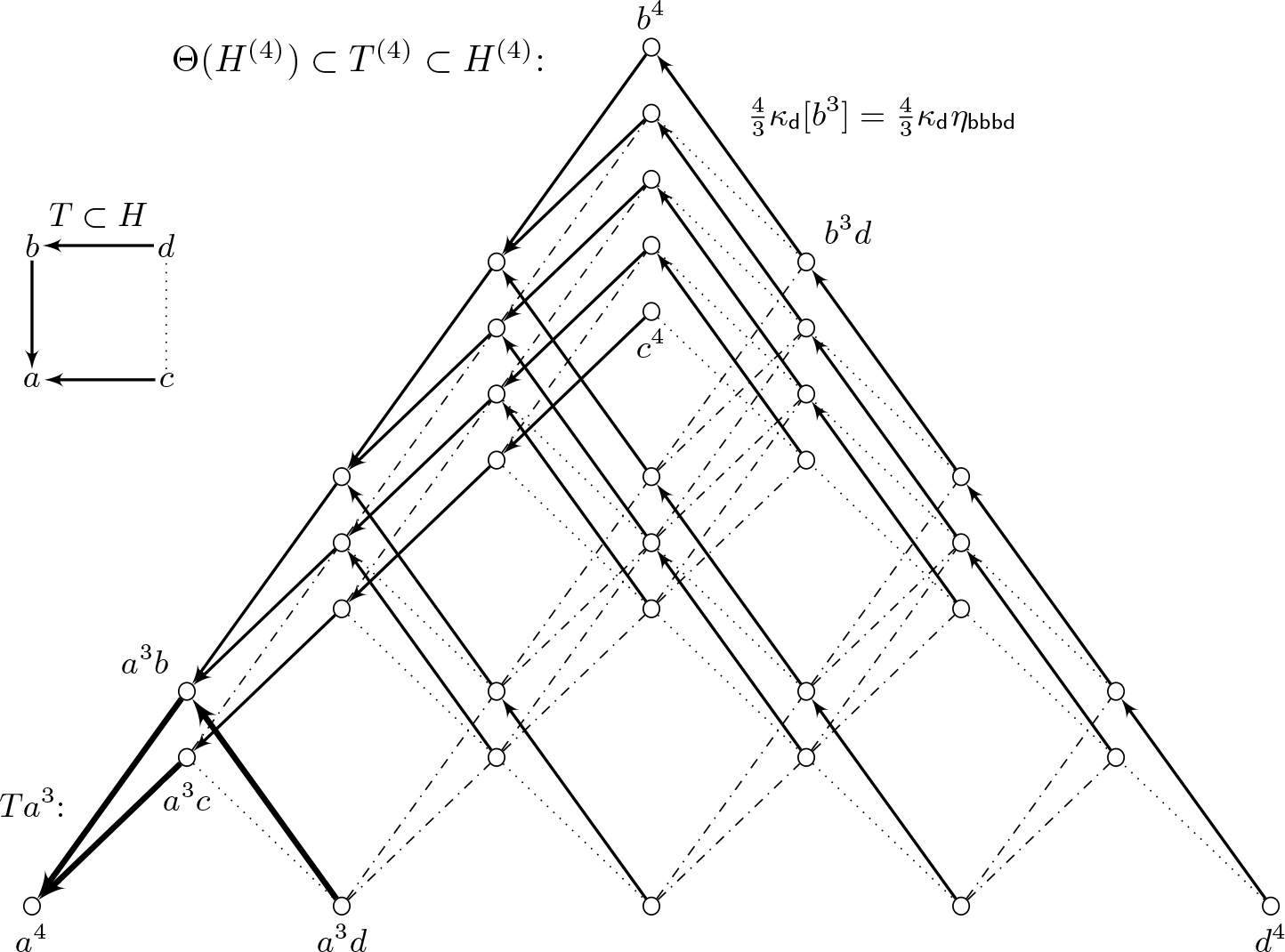
Topology of a receptor oligomer composed of 4 identical ternary complex monomers (*H*, see Fig. 1) is given by the reduced power *H*^(4)^ that contains 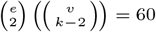 Cartesian squares, but many of these are not independent. The cycle space of *H*^(4)^ has dimension 46. The graph *T*^(4)^ is a subgraph of *H*^(4)^ involving all the edges of *H*^(4)^ that are solid or dash-dotted (but not dotted). The solid arrows are Θ(*G*^*k*^), the subgraph of *T*^(*k*)^ that is the distinguished spanning tree of *G*^(*k*)^

## Supplementary Note S8: Cubical ternary complex dimer

For another example, consider the cubical ternary complex model (denoted *J* in Fig. 7). There are 12 reversible reactions, but only 7 free equilibrium parameters, because *β*(*G*) = *e − v* + 1 = 12 − 8 + 1 = 5. In (49), the 7 free parameters are specified as 3 equilibrium constants (*K_L_*, *K_G_*, and *K_A_* = [R_*a*_]/[R_*i*_]), plus 4 allosteric coupling parameters (*α*, *β*, *γ*, *δ*). Using a spanning tree with R_*i*_ ↔ *a*_1_ as root (Fig. 7, right), we find

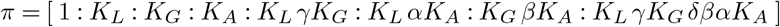

where *π* = [*π*_1_: *π*_2_: ···: *π*_8_] = [[R_*i*_]: [LR_*i*_]: ···: [LR_a_G]].

Fig. S6 shows the structure of cubical ternary complex *homodimer*, denoted by *J*^(2)^, which has 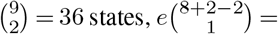 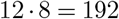 edges (it is not a hypercube), and 192−36+1 = 157 thermodynamic constraints. There are 192−157 = 35 free equilibrium parameters, and the spanning tree Θ(*J*^(2)^) has 35 edges (Fig. 8). The edges removed from *T*(*J*)^(2)^ to create Θ are 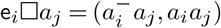 for 2 ≤ *i* < *j* ≤ *v*. These are *e*_2_◻*a*_3_, …, *e*_2_◻*a*_8_, e_3_◻*a*_4_, …, e_3_◻*a*_8_, e_4_◻*a*_5_, …, e_4_◻*a*_8_, e_5_◻*a*_6_, …, e_5_◻*a*_8_, e_6_◻*a*_7_, e_6_◻*a*_8_, e_7_◻*a*_8_. Fig. 8 shows the edges that remain. Seven parameters are inherited from the spanning tree of the monomer model (e_2_, e_3_, …, e_8_ in Fig. 7). The remaining parameters are 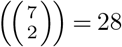 2-way allosteric coupling parameters, denoted 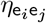 where 2 ≤ *i* ≤ *j* ≤ *v*.

The occupation measure for each state can be found using the token method described in the main text. For example, the term corresponding to *a*_3_*a*_5_ begins with 2*π*_3_*π*_5_. The token diagram is

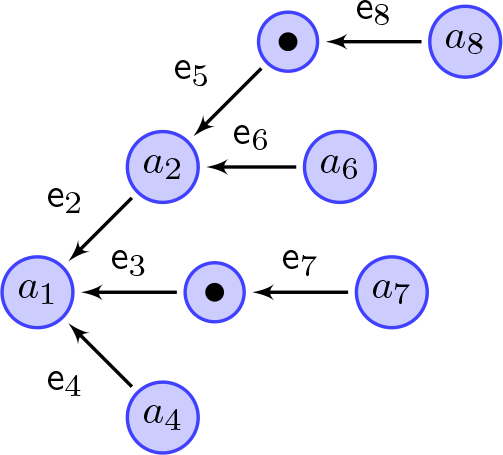

Here *h*_1_ = e_3_ and *h*_2_ = e_2_ + e_5_ (the sum of the edge labels from the token to the root *a*_1_). The 2-way interactions are enumerated by *h*_1_*h*_2_ = e_3_(e_2_ + e_5_) = e_2_e_3_ + e_3_e_5_, so the allosteric factor is 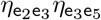. Using 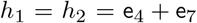, 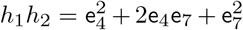, and 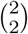 the term for 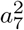 is found to be 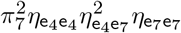. Using a computer algebra system, it is possible automate this procedure to quickly find every term in 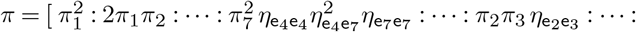 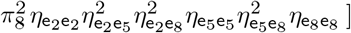 where *π_i_* for the isolated monomer are given above.

## Supplementary Note S9: Automated enumeration of allosteric parameters in receptor oligomers

The enumeration of allosteric parameters in a receptor oligomer can be fully automated. An example Matlab script is provided, along with results corresponding to the example oligomers discussed in the main text.

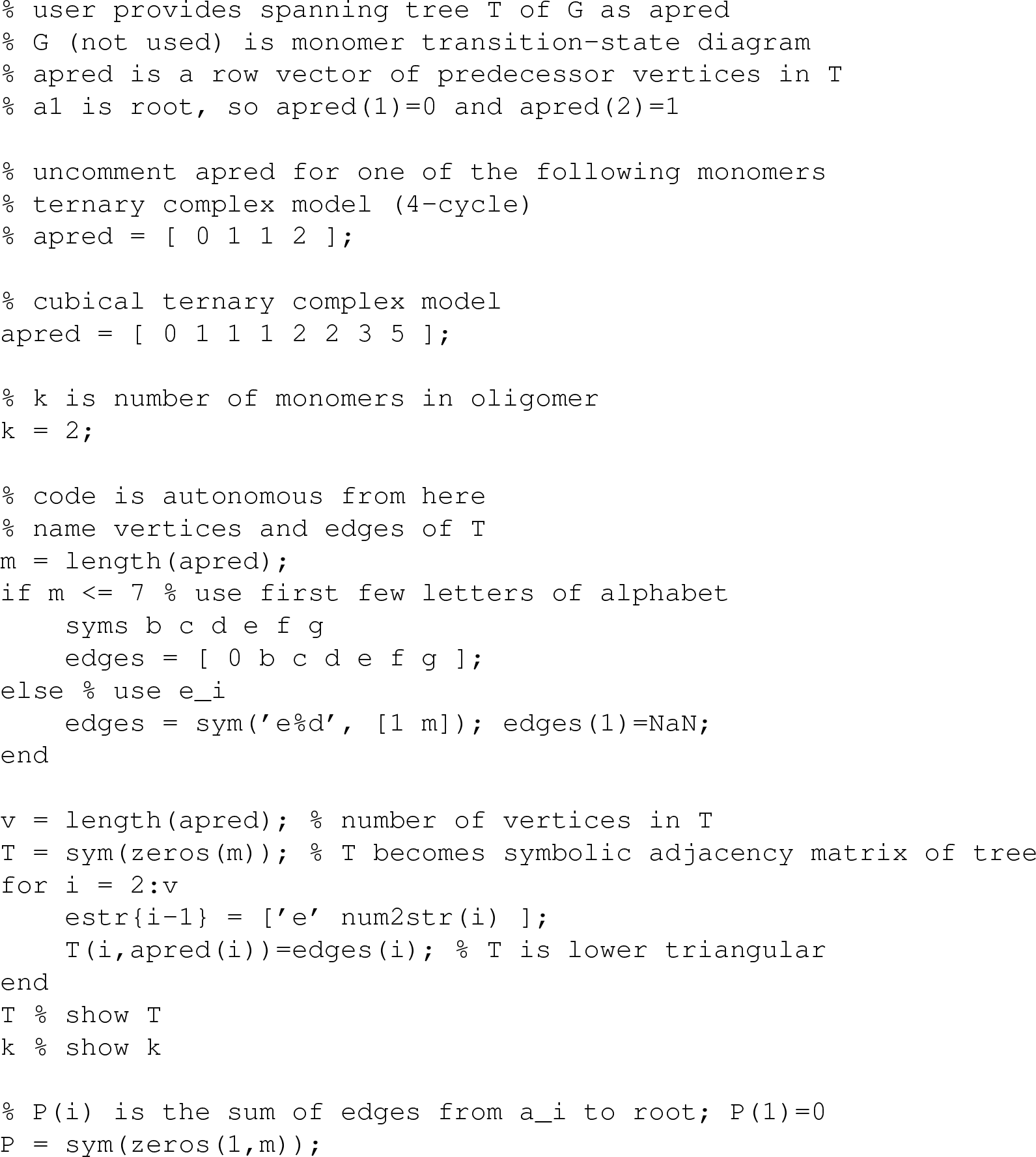

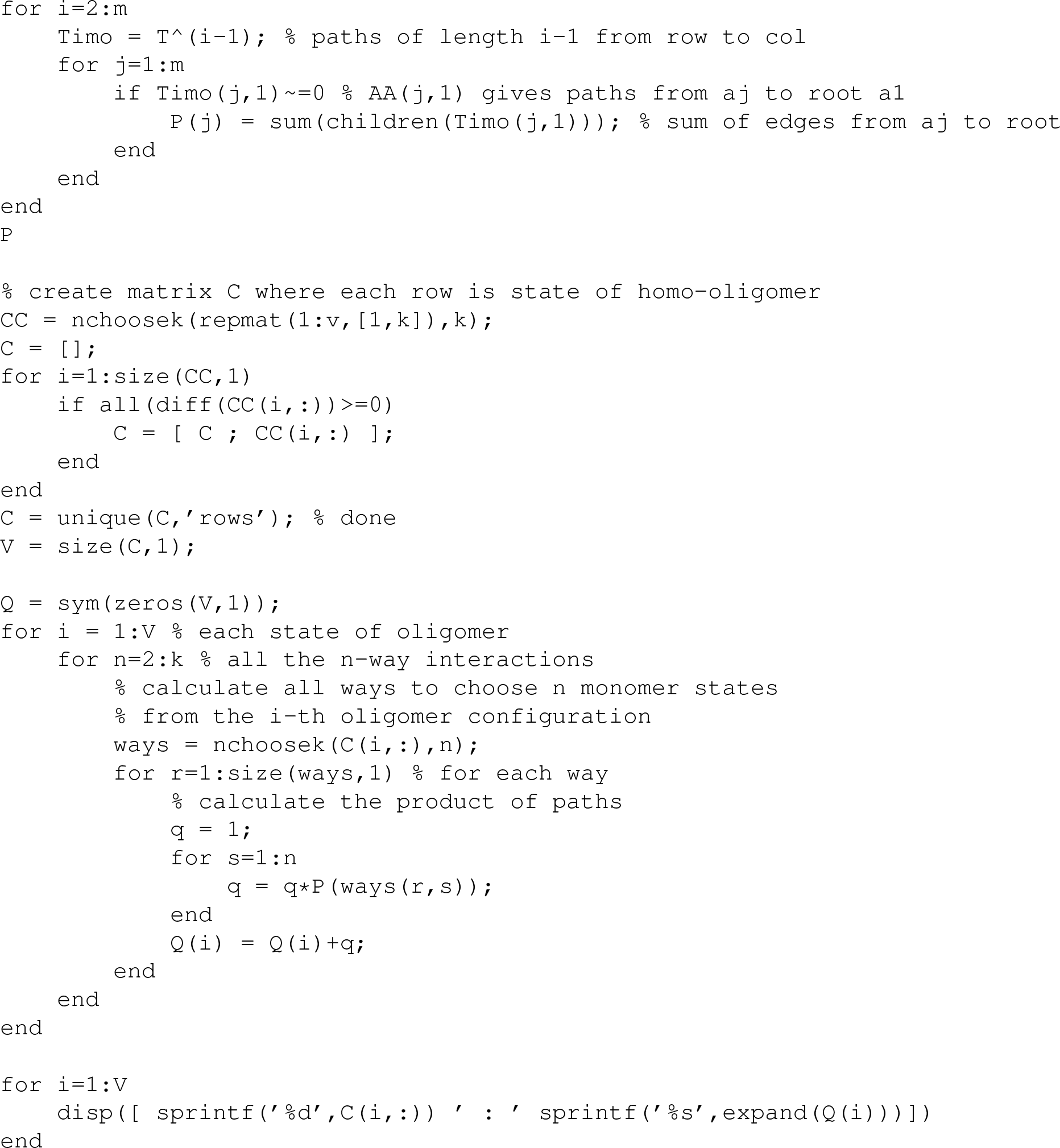

### The ternary complex dimer

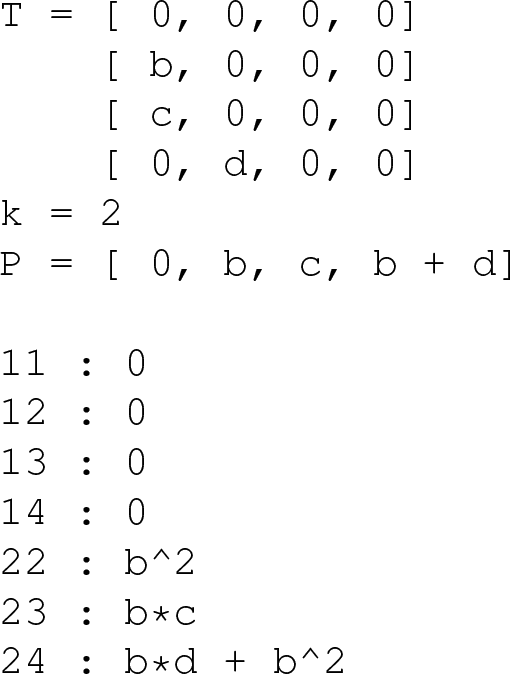

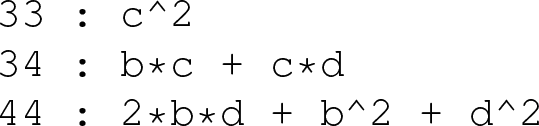

The translation of the script output to the notation used in the main text is straightforward. For example, state 23 means *a*_2_*a*_3_ ~ *bc*, state 44 means 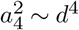), and so on. The allosteric factors for state 23 is *η*_bc_ and for state 44 it is 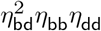, which is written in lexicographical order in the main text: 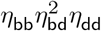.

### The ternary complex tetramer

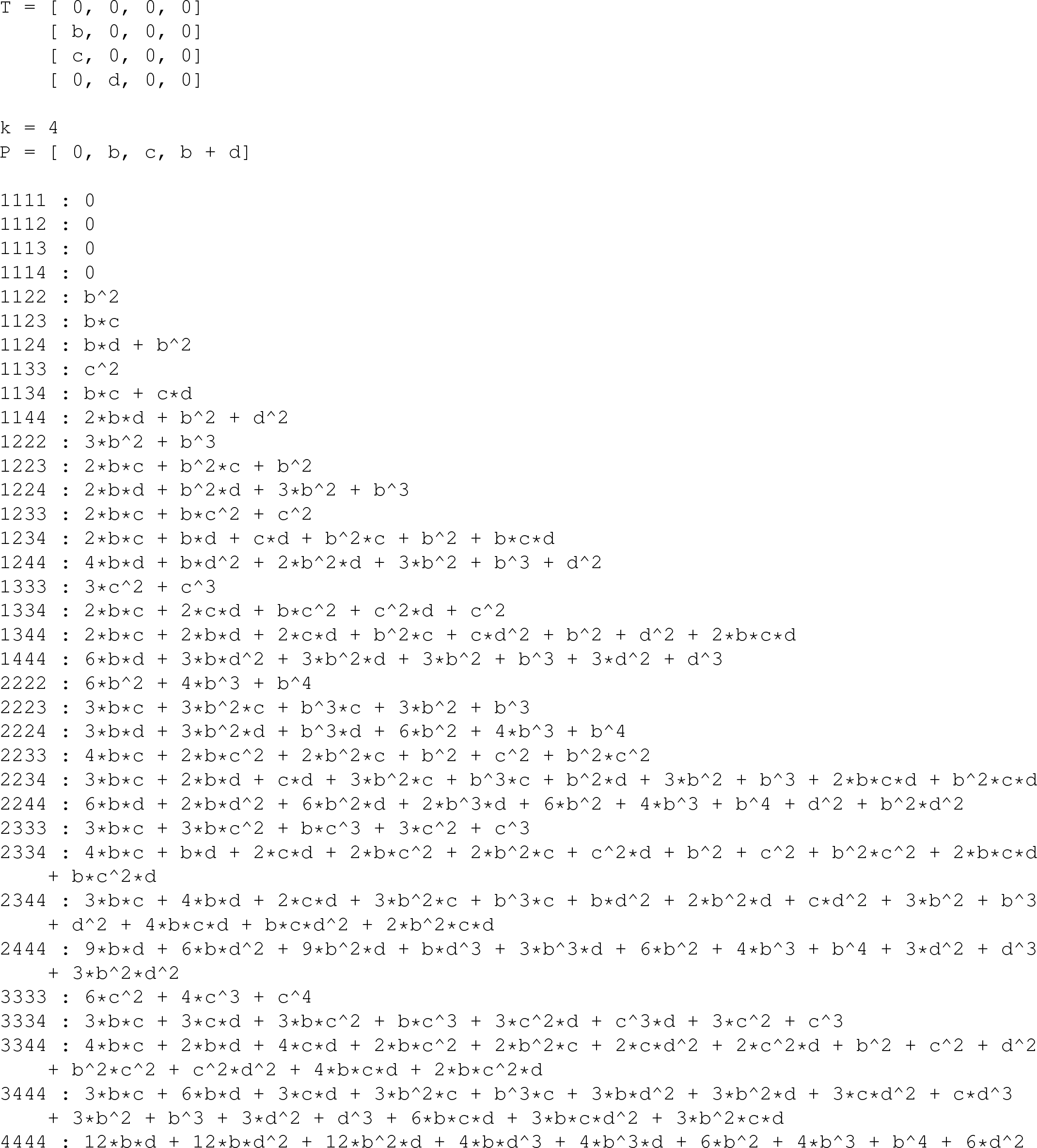

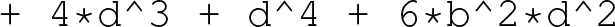

### The cubical ternary complex dimer

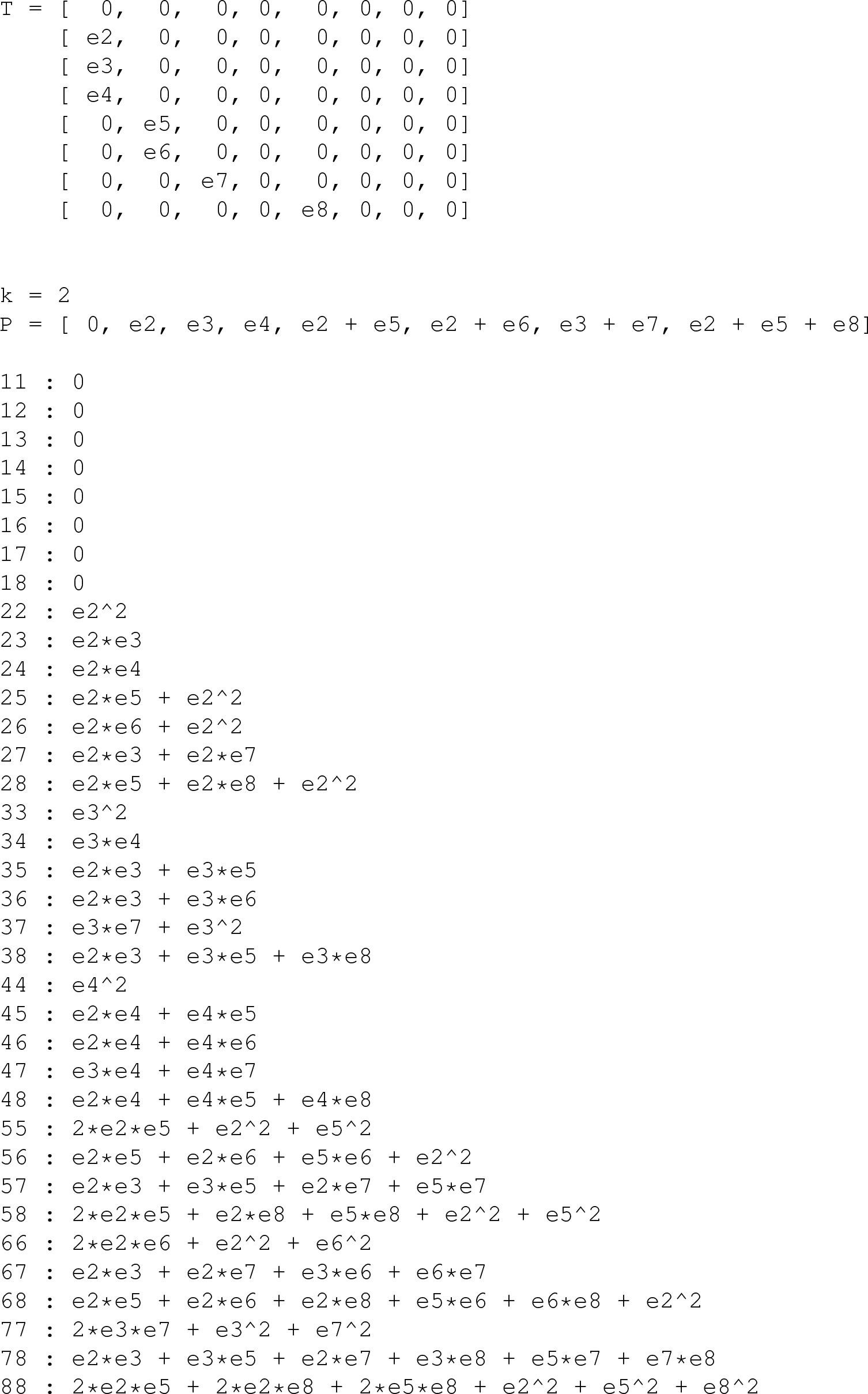

**Fig. S6.**
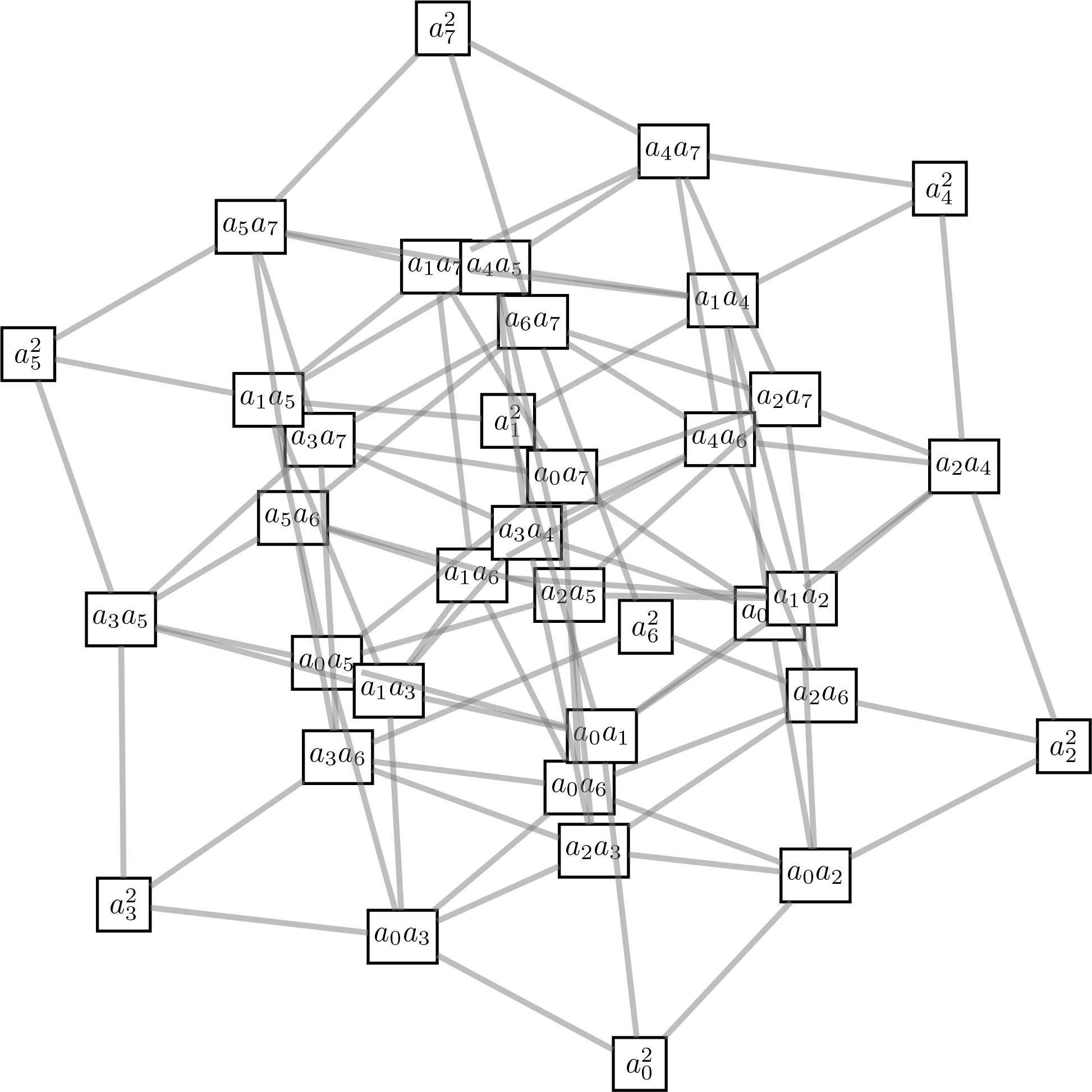
The structure of the cubical ternary complex dimer *J*^(2)^

